# Macrophages drive inguinal fat pad and lymph node remodelling in response to peripheral inflammation

**DOI:** 10.1101/2025.10.06.680668

**Authors:** Robin Bartolini, Deepika Sharma, Gillian J. Wilson, Gillian Dunphy, Jonathan Cavanagh, Jennifer A. Barrie, Heba A. Halawa, John Cole, Stefan Weidt, Kirstyn Gardner-Stephen, Gerard J Graham

## Abstract

Adaptive immune responses are intensely energy-dependent and rely on a local source of fuel-producing molecules which have been proposed to be derived from fat pads in which mammalian lymph nodes are embedded. However, the trigger for their release has not been identified. Here we demonstrate that cutaneous inflammation is directly correlated with rapid atrophy of perinodal fat pads and increase in embedded lymph node size. We further demonstrate that the fat pad atrophy is associated with influx of a CCR2-independent, lipid metabolising, macrophage population and that macrophage depletion ameliorates fat pad atrophy, and lymph node expansion, downstream of inflamed sites. Our data therefore identify peripheral inflammation as a trigger of downstream fat pad and lymph node remodelling and contributes to the release of essential nutrients to drive the energetic requirements of the adaptive immune response.

**Significance statement:** To our knowledge, nobody has previously reported this striking correlation between peripheral inflammation and reciprocal fat pad and lymph node remodelling. Our report of this correlation and our mechanistic results, have clear implications for our understanding of the inflammation-driven rapid release of sources of energy from fat pads to drive the immune response. Our data also potentially shed light on additional aspects of the functionality of pro-inflammatory vaccine adjuvants. We believe that our findings are fully novel and will be of interest to immunologists, infectious disease specialists and researchers interested in adipose tissue derived energetics.

## INTRODUCTION

In adaptive immune responses, antigen presentation from peripheral tissues via dendritic cells triggers specific lymphocyte activation with the generation of target-specific T cell responses and ultimately the generation of immune memory(1). Importantly, these events must take place within the context of an inflammatory, or damage-related, challenge at the site of antigen encounter(2). There is also ample evidence of other peripheral tissue-derived molecules providing additional specificity to the immune response especially in terms of assigning selective tissue-homing properties to the antigen-specific T cells(3, 4). This is therefore a complex response which requires the simultaneous, and coordinated, interaction of multiple cell types and molecules. In mammals this is achieved by integrating all these processes within lymph nodes that lie downstream of all tissues and which are connected to the tissues via the lymphatic vasculature(5).

In addition to requiring precise coordination of cells and molecules, the immune response is highly energy-intensive and requires rapid access to local molecular energy sources including free fatty acids(6). Indeed, research indicates that there is a trade-off between the high metabolic costs of immune activation and the maintenance of body temperature(7). In this regard, it is notable that all mammalian lymph nodes are embedded in perinodal adipose tissue(8) which contains, not just adipocytes, but also a range of immune cells with macrophages representing the largest resident population. Considerable research indicates a key role for the perinodal adipose tissue as a source of nutrients, released through lipolysis, to feed the immune response(9). In addition, there is clear evidence of an effect of immune stimulation on perinodal adipose tissue content and size(10). What is not understood is how this process linking adipocytes in perinodal adipose tissue to immune response is regulated.

Here we demonstrate that induction of cutaneous inflammation is sufficient, in the absence of antigen presentation, to remodel lymph nodes and perinodal fat deposits. Specifically cutaneous inflammation induced either by wounding, or administration of the TLR9 agonist imiquimod, is associated with rapid atrophy of the fat pad embedding the inguinal lymph node and a marked increase in size of the lymph node. Both effects are strongly correlated with the degree of peripheral inflammation and are dependent on the CCR2-independent accumulation of a specific macrophage population in the fat pad. We therefore identify inflammation as a trigger for perinodal fat pad, and lymph node, remodelling in the immune response. These data therefore shed new light on the communication between peripheral tissues and lymph nodes in the initiation of an immune response and suggest that the mechanisms of action of pro-inflammatory vaccine adjuvants may not be restricted to the peripheral injection site.

## RESULTS

### CCR2−/− male mice display reduced inguinal fat pad, and increased inguinal, lymph node size

Gross examination of resting male CCR2−/− mice revealed a striking phenotype in the inguinal fat pad, characterised by a marked reduction in the size of the fat pad and a concomitant increase in the size of the embedded inguinal lymph node (Figure 1A). Quantitative analyses confirmed these observations with a significant reduction in inguinal fat pad size and weight (Figure 1Bi and ii) and an increase in inguinal lymph node volume and cellularity (Figure 1Ci and ii). Overall, a strong negative correlation between lymph node weight and fat pad weight was observed in CCR2−/− males (Figure 1D). Importantly this phenotype was only observed in male mice (Figure 1A) and progressed with age (Figures 1B and 1C). Total body weight was unaffected in male (and female) mice (Figure 1E), suggesting that the phenotype observed in male mice is not systemic.

**Figure 1.**
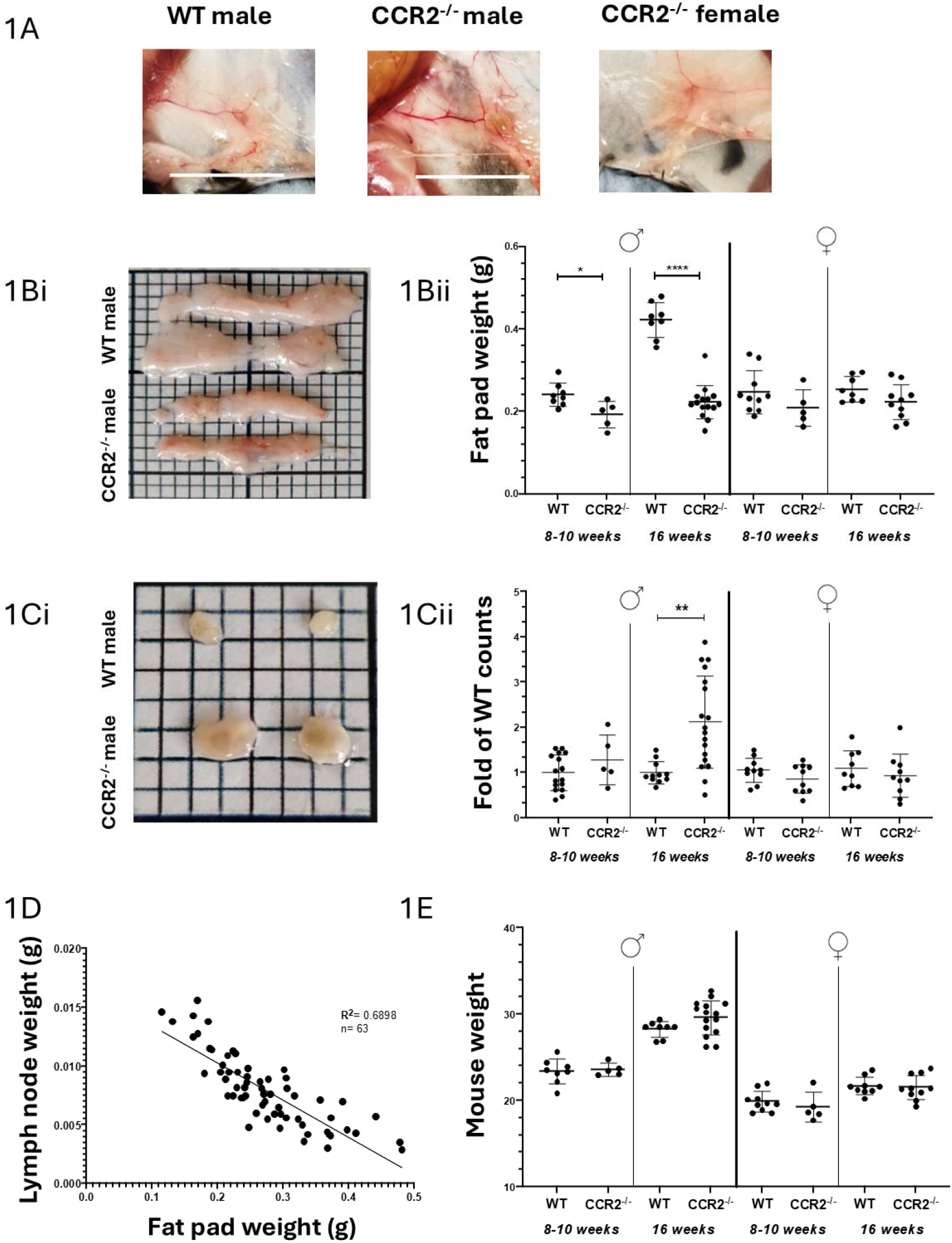
CCR2−/− male mice display reduced inguinal fat pad, and increased inguinal, lymph node size. **A**: In situ orientation of inguinal fat pads with embedded inguinal lymph nodes in WT male (C57BL/6 N), CCR2−/− male and CCR2−/− female mice. Scale bar: 1cm. **B**: i) Excised inguinal fat pads from WT male (top) and CCR2−/− male (bottom). Scale bar: 1cm. ii) Inguinal fat pad weight (in grams) of WT and CCR2−/− mice, both male and female, at 8-10 weeks and 16+ weeks of age. Each point represents the combined weight of both inguinal fat pads of a single mouse. **C**: i) Excised inguinal lymph nodes from WT male (top) and CCR2−/− male (bottom). Scale bar: 0.5cm. ii) Live cell counts of inguinal lymph nodes, expressed as fold of WT mice, both male and female, at 8-10 weeks and 16+ weeks of age. Each point represents the combined weight of both inguinal lymph nodes of a single mouse. The number of live cells was determined via dye exclusion of trypan blue and normalised to the average of WT counts (age and gender matched) to obtain values expressed as fold of WT. **D:** Correlation of inguinal fat pad (x-axis) and lymph node (y-axis) weight, both expressed in grams, showing a considerable degree of dispersion and variability at the resting state in CCR2^−/−^ males. Each point corresponds to the combined weight of both fat pads and both lymph nodes of a single CCR2−/− mouse. Simple linear regression analysis was conducted, yielding an equation of Y=-0.03174x+0.01662 (plotted, black), with an R^2^ value of 0.6898. The slope coefficient was found to be significantly non-zero, as indicated by a p value of <0.0001. N= 63 **1E**: Overall mouse weight, in grams, of WT and CCR2^−/−^ mice, both male and female, at 8-10 weeks and 16+ weeks of age. Unpaired t test with Welch’s correction was performed to determine statistical significance, with a p value of 0.05 determined as significant. *p<0.05, **p<0.01, ****p<0.0001

Thus, CCR2−/− male mice display reduced inguinal fat pad weight and increased inguinal lymph node size which are correlatively linked.

### The observed phenotype is associated with skin wounding

The fact that the above differences were observed in male, but not female, mice suggested a possible association of fighting-induced inflammation or infection to the phenotype. Therefore, we examined the contribution of co-housing to the phenotype in CCR2−/− male mice. To this end, CCR2−/− males (8 weeks old) were either segregated, or co-housed in cages of 5, and culled 3 weeks later. As expected, co-housed CCR2−/− male mice showed evidence of more severe, fighting-related, skin wounding than their segregated counterparts. Of note, some co-housed CCR2−/− male mice had to be culled due to excessive wounding (Figure 2A). This co-housing was clearly associated with smaller inguinal fat pads and larger inguinal lymph nodes in comparison to segregated counterparts (Figures 2B-D). As noted above, we observed a strong negative correlation between lymph node weight and fat pad size in CCR2−/− males (Figure 1D). When separating co-housed and segregated CCR2−/− males, we obtained 2 different functions: segregated CCR2−/− males showed no significant correlation between lymph node weight and fat pad size (Figure 2Ei), which was similar to what was observed in WT males (Figure 2Eii) and CCR2−/− females (Figure 2Eiii). In contrast, co-housed CCR2−/− males showed a very strong correlation between lymph node and fat pad weight (Figure 2Eiv), indicating that this correlation only holds for co-housed CCR2−/− mice (Figure 2F).

**Figure 2.**
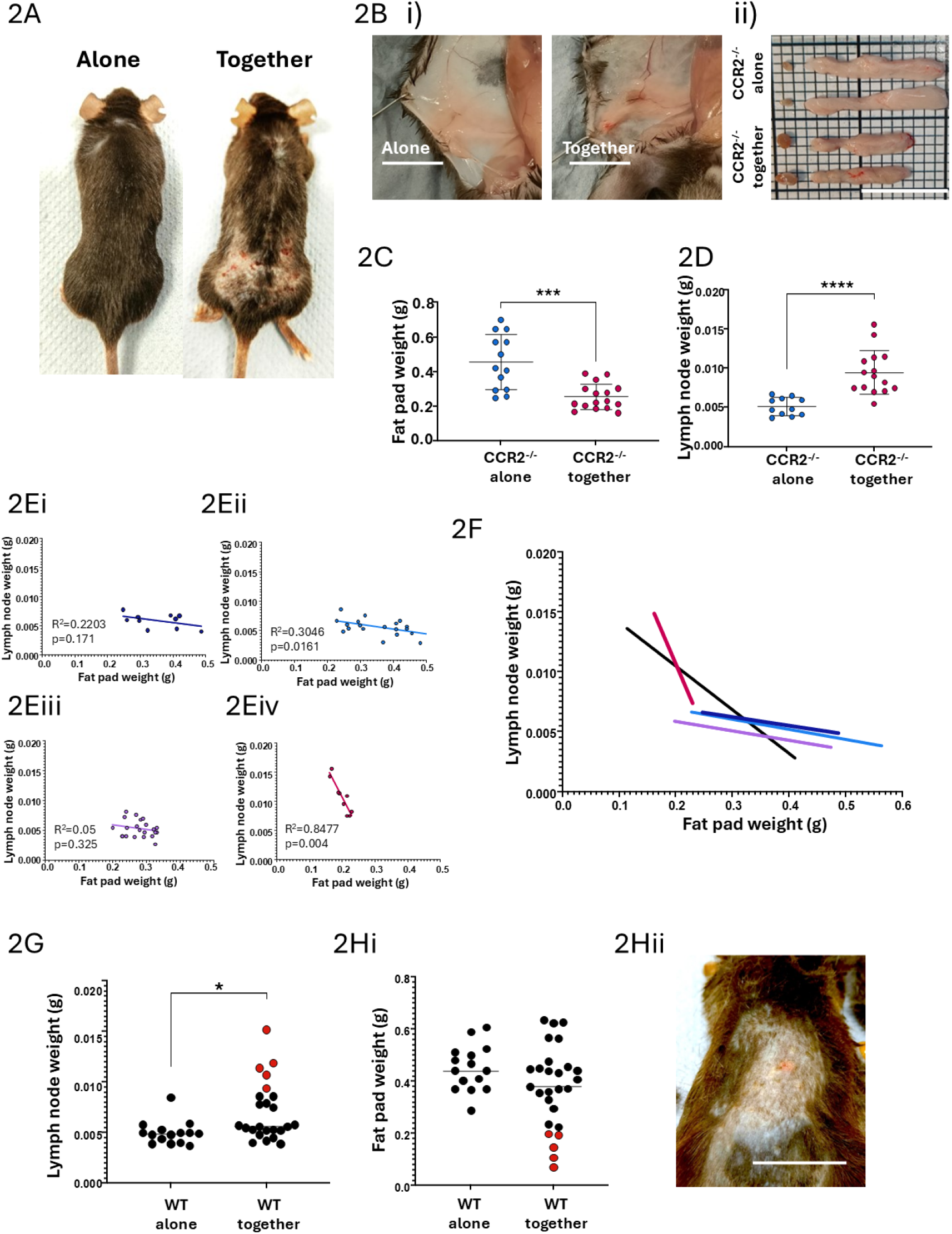
The observed phenotype is associated with skin wounding. **A**: Examples of more severe fighting-induced injuries observed in CCR2−/− males housed together in groups of 5 from the age of 8 weeks compared to CCR2−/− males housed alone. **B**: i) In situ and ii) excised (top=alone, bottom=together) inguinal fat pads and lymph nodes from CCR2−/− males housed alone or together in groups of 5. Scale bar= 1cm **C**: Inguinal fat pad weight, in grams, of 14-16 week old CCR2^−/−^ mice, either housed together in groups of 5 or alone, from the age of 8 weeks. Both inguinal fat pads of each mouse are pooled for each individual measurement. **D**: Inguinal lymph node weight, in grams, of 14-16 week old CCR2−/− mice, either housed together in groups of 5 or alone, from the age of 8 weeks. Both inguinal lymph nodes of each mouse are pooled for each individual measurement. **E**: Correlation of inguinal fat pad (x-axis) and lymph node (y-axis) weight, both expressed in grams, of CCR2−/− males housed alone (i: dark blue), WT males (ii: light blue), CCR2−/− females (iii: purple) and CCR2−/− males housed together (iv: red). Simple linear regression for CCR2−/− alone, WT and CCR2−/− females was non-significant, with R2 values of 0.2203, 0.3046 and 0.05 respectively. The slope coefficient of the simple linear regression on CCR2^−/−^ housed together was significantly non-zero (p=0.004), with an R^2^ value of 0.8477. **F**: Combined linear regressions of the correlation between Lymph node weight and fat pad weight of CCR2−/− housed alone (blue), CCR2−/− housed together (red), WT (blue), CCR2−/− females (purple) and previously calculated linear regression on 63 CCR2−/− males (black, Figure 1D) **G**: Inguinal lymph node weight, in grams, of WT males at 14 weeks housed either alone or together in groups of 5 from the age of 8 weeks. Each dot represents the combined weight of both inguinal lymph nodes of each mouse. Red dots indicate mice with obvious signs of fighting injuries. **H**: i) Fat pad weight, in grams, of WT males at 14 weeks housed either alone or in groups of 5 from the age of 8 weeks. Each dot represents the combined weight of both inguinal fat pads of each mouse. Red dots indicate mice with obvious signs of fighting injuries. ii) Representative image of fighting induced skin lesions observed in WT males (one of the mice marked as red dot in graphs 2G and 2Hi) housed together in groups of 5 per cage. Scale bar=2cm Unpaired t test with Welch’s correction was performed in 2C-E and 2G and H to determine statistical significance, with a p value of 0.05 determined as significant. *p<0.05, ***p<0.001, ****p<0.0001.

Whilst the above phenotype was not immediately apparent in WT males, subsequent analysis comparing isolated, and co-housed, WT males revealed a small, but significant, increase in lymph node size, and a reduction in fat pad size, in co-housed WT male mice (Figure 2G and 2Hi). These differences were the consequence of a few males in the co-housed population with markedly enlarged lymph nodes and atrophied fat pads (red dots in Figures 2G and Hi). Interestingly, the mice with the most extensive alterations in lymph node and fat pad weight, also showed evidence of fighting-associated skin wounds (Figure 2Hii) thus further suggesting a link between the observed phenotypes and cutaneous inflammation.

Thus, these data link fighting-associated skin inflammation to inguinal fat pad and lymph node alterations.

### CCR2−/− mice display extensive wounding-related skin inflammation

Next, we examined why this phenotype was particularly prominent in co-housed CCR2−/− male mice. Counterintuitively, and despite the fact that CCR2−/− mice display no alterations in myeloid or lymphoid populations apart from profound monocytopenia(11, 12), co-housed CCR2−/− males had significantly more CD45+ cells in the skin (Figure 3A) than segregated CCR2−/− mice, or co-housed, WT mice. This was reflected in a significant increase in neutrophils, eosinophils and T-cells (Figures 3B and 3C). Despite the anticipated reduction in infiltrating monocytes in the co-housed CCR2−/− mouse skins compared to wild-type mouse skins, the co-housed CCR2−/− mouse skins showed significantly more inflammatory monocyte content than single housed CCR2−/− mice suggesting CCR2-independent monocyte recruitment. (11, 12) (Figure 3Di). Strikingly, macrophage numbers were similar in WT and co-housed CCR2−/− mice (Figure 3Dii) with the macrophages expressing high levels of CD11b again reflecting likely derivation from recruited cells rather than local proliferation(13). Further staining revealed the presence of both CD206+ and CD206- macrophages in WT skin, but only CD206+ macrophages in co-housed CCR2−/− mouse skin (Figure 3Diii). Importantly, the number of CD45+ cells in the skin displayed a strong negative correlation with fat pad weight (Figure 3E), indicating that the extent of cutaneous inflammation is linked to inguinal lymph node and fat pad size.

**Figure 3.**
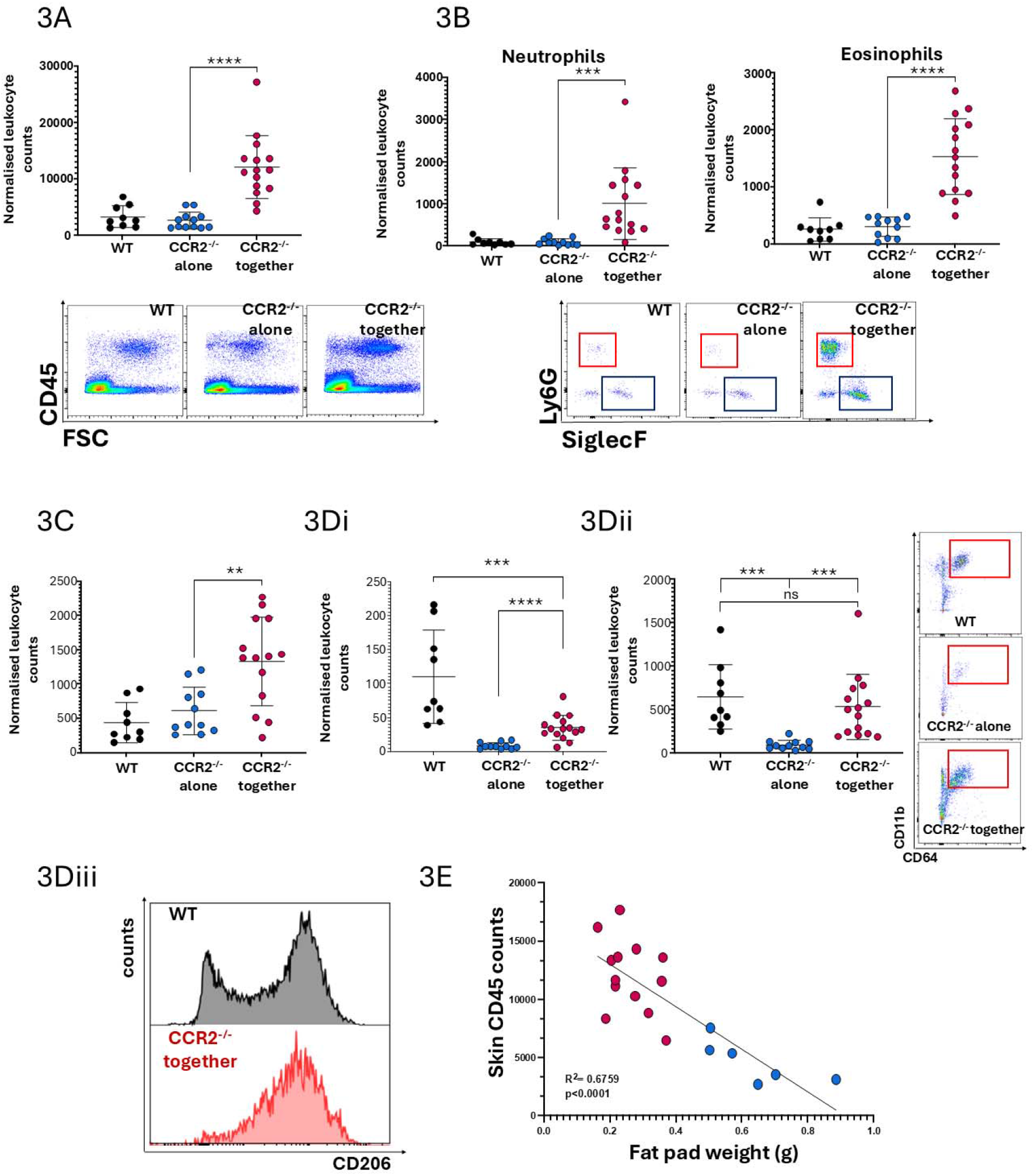
CCR2−/− mice display extensive wounding-related skin inflammation. **A**: CD45+ cell content of the skin, assessed via flow cytometry, in WT (black), CCR2−/− housed alone (blue) and CCR2^−/−^ housed together (red) male mice, with representative FACS plots for each below. **B**: Neutrophil and eosinophil numbers in the skin of WT (black), CCR2^−/−^ housed alone (blue) and CCR2−/− housed together (red) males, with representative FACS plots for each below (neutrophils= CD11b+Ly6G+ red box, eosinophils= CD11b+SiglecF+ blue box). **C**: T-cell numbers (CD11b+ CD3+ CD19-) in the skin of WT (black), CCR2−/− housed alone (blue) and CCR2−/− housed together (red) male mice **D**: i) Inflammatory monocyte (CD11b+Ly6C++ Ly6G-) and ii) macrophage (CD11b+ CD64+) numbers in the skin of WT (black), CCR2^−/−^ housed alone (blue) and CCR2−/− housed together (red) males, with representative FACS plots for CD11b/CD64 staining highlighting macrophages (red box). iii) Representative histograms showing CD206 staining, assessed via flow cytometry, on the CD11b+CD64+ macrophage population in the skin of WT males (black histogram) and CCR2−/− males housed together (red histogram) **3E**: Correlation of fat pad weight (in grams, x-axis) and normalised CD45+ cell counts in the skin of CCR2−/− males (y-axis), alone (blue dots) and housed together (red dots). Fat pad weight values represent the pooled weight of both fat pads for each mouse. Simple linear regression analysis was conducted (black line), with an R^2^ value of 0.6759 and a slope coefficient significantly non-zero, as indicated by a p value of <0.0001. N=19 (6 resting, 13 inflamed) For 3A-D, Leukocyte numbers were normalised to the weight of the skin sampled. Unpaired t test with Welch’s correction was performed to determine statistical significance in 3A-D with a p value of 0.05 determined as significant. **p<0.01 ***p<0.001, ****p<0.0001

Thus CCR2−/− mice display exaggerated skin inflammation, the extent of which correlates with the reduction in inguinal fat pad weight.

### Inguinal fat pads and lymph nodes in CCR2−/− mice display altered cellularity

Next, we examined the leukocyte content of the remodelled fat pads and lymph nodes. Whilst, as expected, fat pads from co-housed male CCR2−/− mice were severely depleted in inflammatory monocytes (Figure 4Ai), they displayed a general increase in macrophage numbers (Figure 4Ai and ii) compared to WT mice. Further flow cytometric analysis for expression of CD11c, a discriminating marker in adipose tissue macrophages(14) and a broad marker of inflammatory macrophages(15), revealed an increased number of pro-inflammatory CD11c+ CD206+ macrophages in CCR2−/−, compared to WT, male mice and a significant reduction in CD11c-CD206+ anti-inflammatory macrophages (Figure 4Bi, ii and iii). We suggest that these cells are equivalent to the recently described F4/80+TIM4-CD11c+ population of white adipose tissue macrophages(14). Again, these phenotypes progressed with age and were not seen in female CCR2−/− mice. All other tested fat pad depots in CCR2−/− males and females showed a significant reduction in the number of infiltrating inflammatory monocytes with a concomitant decrease in the number of adipose tissue macrophages (ATMs) in visceral and brown fat but not in axillary, fat depots (Supplementary Figure 1A). These data suggest that the alterations in fat pad monocyte and macrophage numbers is largely restricted to fat pads downstream of the inflamed skin and is therefore likely to be a consequence of the fighting-related cutaneous inflammation. Further characterisation of the inguinal ATM in CCR2−/− male mice revealed increased expression of CD11b and other inflammatory markers (Supplementary Figure 1B and 1C). As with macrophages in the inflamed skin, the expression of CD11b and CD11c on the fat pad macrophages is consistent with recruitment, again suggesting CCR2-independent mechanisms. In addition to macrophages, flow cytometric analysis revealed a significant increase in both neutrophils and eosinophils in the inguinal fat pads of male CCR2−/− mice which, again, increased with age (Figures 4Ci, 4Cii, 4Ciii) and was not seen in female mice. Importantly, the increased numbers of macrophages, neutrophils and eosinophils was significantly ameliorated when the CCR2−/− male mice were isolated from each other again confirming the role for fighting in this phenotype (Supplementary Figure 2A). Analysis of visceral fat showed no differences between co-housed and segregated CCR2−/− male mice (Supplemental Figure 2B), with normal numbers of leukocytes (Supplemental Figure 2C), but a reduction in monocytes and macrophages compared to WT (Supplemental Figure 2D).

**Figure 4.**
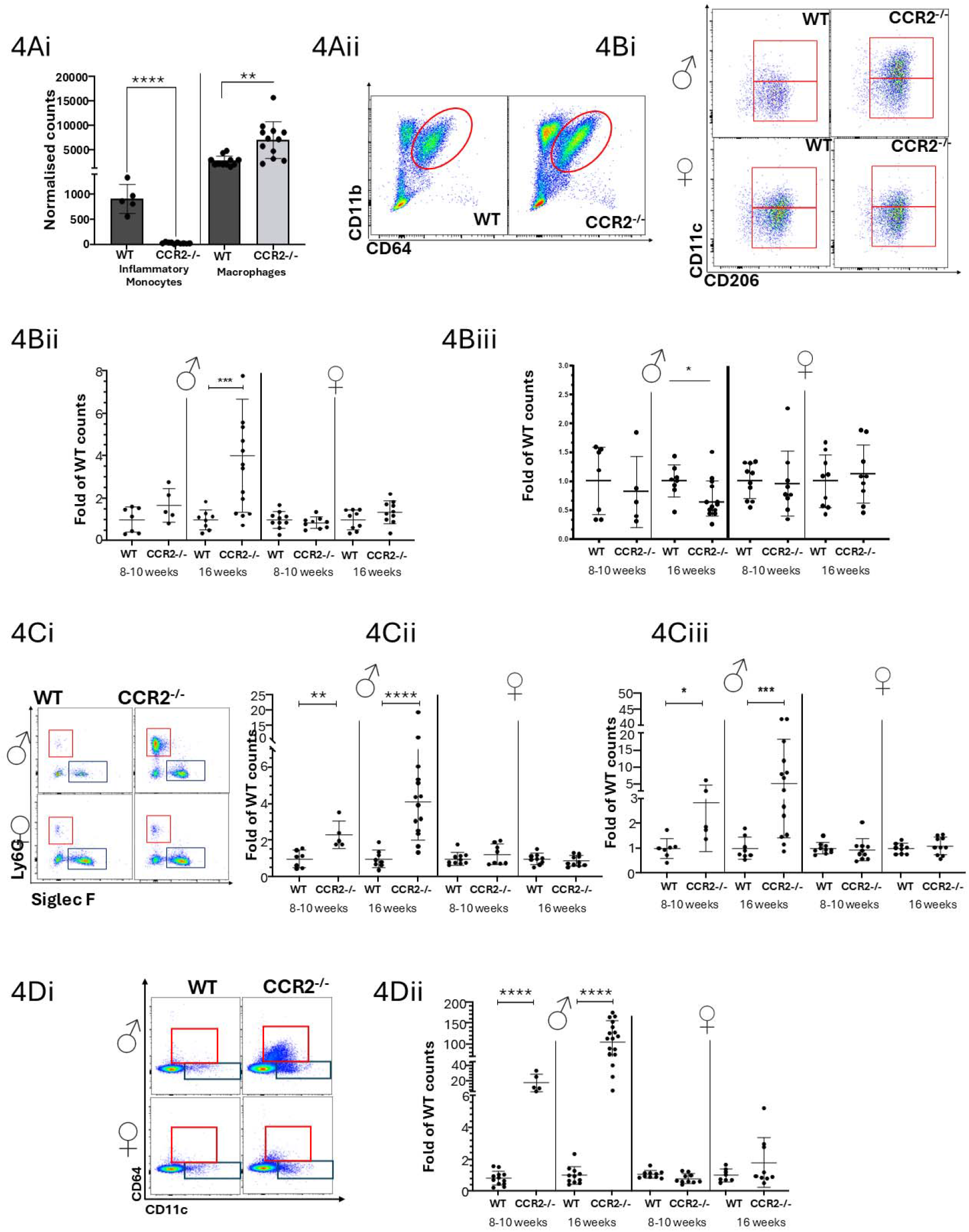
Inguinal fat pads and lymph nodes in CCR2^−/−^ mice display altered cellularity. **A**: i) Inguinal Fat pad infiltrating inflammatory monocyte (CD11b+CD64-Ly6C++Ly6G-) and macrophage (CD11b+CD64+) normalised counts in WT and CCR2−/− males. Both inguinal fat pads for each mouse were pooled to obtain one measurement, and counts were normalised to the combined weight of both fat pads. ii) Representative FACS plots showing CD11b+ CD64+ macrophages in the inguinal fat pads of WT and CCR2−/− male mice (red oval). **B**: i) Representative FACS plots showing further CD11c/CD206 staining of the CD11b+CD64+ inguinal fat pad macrophage population found in WT and CCR2^−/−^ mice, both male (upper panels) and females (lower panels). ii) Number of CD11c+ and iii) CD11c- adipose tissue macrophages in WT and CCR2−/− inguinal fat pads, in both males and females, at 8-10 weeks and 16+ weeks of age. Each point represents the combined macrophage content of both inguinal fat pads of a single mouse. **C**: i) Representative FACS plots showing CD11b+ CD64- neutrophils (Ly6G+, red box) and eosinophils (SiglecF+, blue box) in the inguinal fat pads of WT and CCR2−/− mice, both male and female. Quantification of inguinal fat pad ii) eosinophils and iii) neutrophils in WT and CCR2−/− inguinal fat pads, in both males and females, at 8-10 weeks and 16+ weeks of age. Each point represents the combined cellular content of both inguinal fat pads of a single mouse. **D**: i) Representative FACS plots showing subcapsular macrophages (CD3- CD19- CD64+ CD11c-, red box) and dendritic cell (CD3- CD19- CD64- CD11c+, blue box) and ii) quantification of subcapsular macrophage numbers in the inguinal lymph node of WT and CCR2−/− mice, both male and female, at 8-10 weeks and 16+ weeks of age. Each point represents counts from both inguinal lymph nodes for each mouse. For 4B-D Values were normalised to the average of WT counts (age and gender matched) to obtain values expressed as fold of WT. Unpaired t test with Welch’s correction was performed to determine statistical significance in 4Ai, 4C-D, with a p value of 0.05 determined as significant. *p<0.05, **p<0.01, ***p<0.001, ****p<0.0001.

In the inguinal lymph nodes of CCR2−/− male mice, we observed a 20-fold increase (8-10 weeks) and up to 150-fold increase (16 weeks) in numbers of subcapsular macrophages (Figure 4Di and ii). This increased lymph node cellularity was not restricted to macrophages and whilst no differences were observed at 8-10 weeks of age, inguinal lymph nodes of CCR2−/− males displayed increased numbers of B-cells and CD4 and CD8 T-cells by 16 weeks of age (Supplementary Figure 3A). Again, female mice did not show this phenotype. Mandibular and mesenteric lymph nodes showed no increase in subcapsular macrophage numbers between WT and CCR2−/− males (Supplementary Figure 3B), suggesting that the increased lymph node size is restricted to the inguinal lymph nodes.

Thus, both the atrophied fat pad, and enlarged lymph node, are characterised by increased levels of resident leukocytes.

### Reduced inguinal fat pad weight and increased inguinal lymph node weight is independent of iCCRs

The data above indicate emergence of an ATM population in the fat pads of male CCR2−/− mice despite severe depletion of inflammatory monocytes. We therefore next investigated roles for other myeloid-related chemokine receptors (inflammatory CCRs: iCCRs) in ATM recruitment by analysing the leukocyte content of the inguinal fat pad of iCCR−/−(11) (compound deletion of CCR1, CCR2, CCR3, and CCR5) male mice. Importantly, these mice phenocopy CCR2−/− male mice in that they display equivalent reciprocal changes in fat pad and lymph node weights (Figure 5Ai and ii). Flow cytometry analysis indicated that, like CCR2−/− mice, the iCCR−/− male fat pads displayed a generally increased inflammatory cellular content with a particularly significant increase in neutrophils (Figure 5Bi and ii). In addition, an increase in CD11c+ CD206+ ATMs was also seen (Figure 5C) which increased as a proportion of total ATMs as mice aged (Figure 5D). Importantly, numbers of eosinophils in the iCCR−/− fat pads were not increased likely as a direct result of the deletion of the eosinophil receptor CCR3 (Figure 5Bii). Thus, these data show that other iCCRs do not contribute to the fat pad macrophage accumulation seen in CCR2−/− mice and discount eosinophils as a cell type contributing to the phenotype.

**Figure 5.**
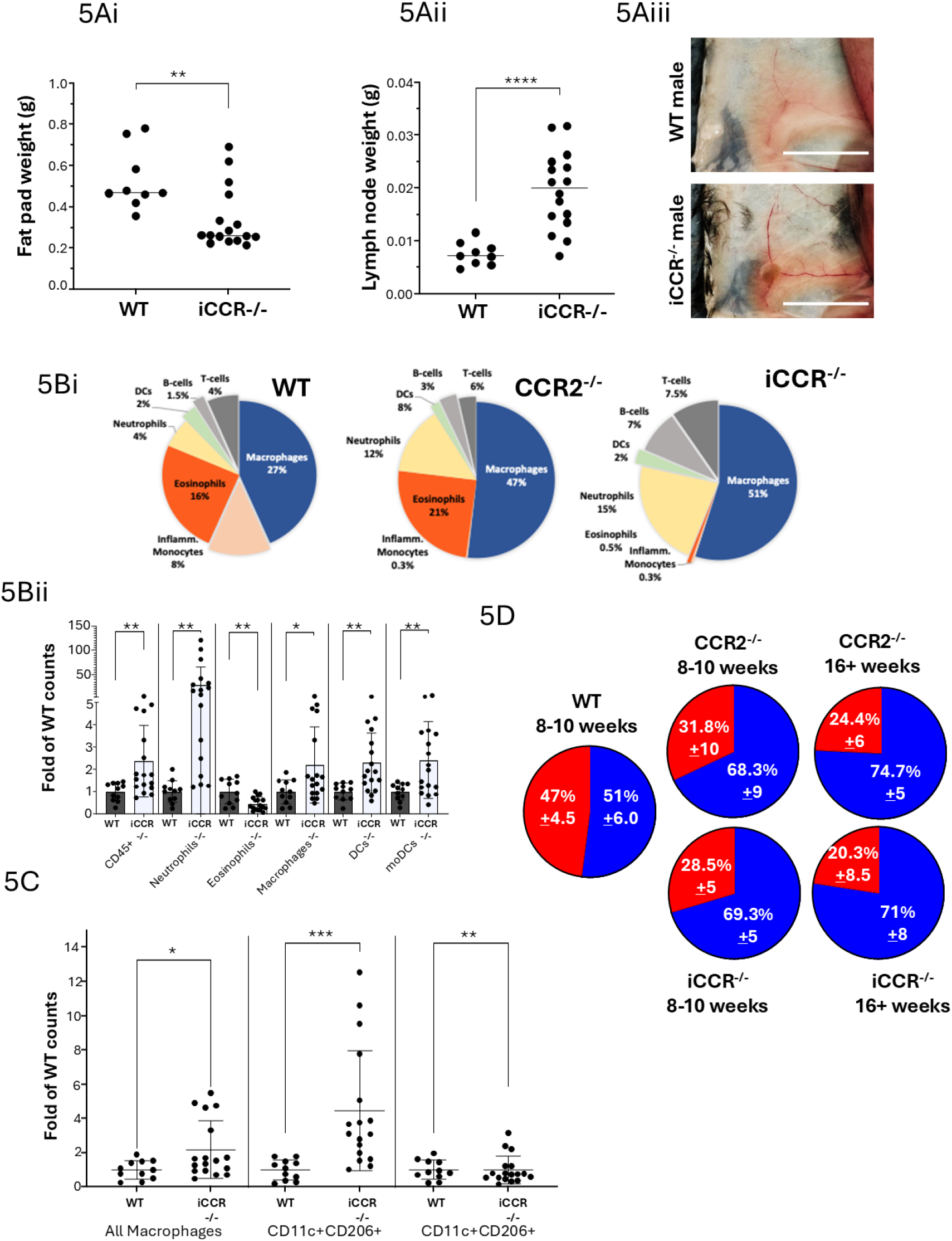
Reduced inguinal fat pad weight and increased inguinal lymph node weight is independent of iCCRs. **A**: i) Inguinal fat pad weight and ii) inguinal lymph node weight, in grams, of 14-16 weeks old male iCCR−/− mice compared to age and gender matched WT mice (with representative images (iii) showing in situ orientation for both, WT mice (top panel) and iCCR−/− mice (bottom panel), scale bar=0.5cm). Both inguinal fat pads and inguinal lymph nodes of each mouse are pooled for each individual measurement. **B**: i) leukocyte subset frequencies in the inguinal fat pad of male mice of 16 weeks of age in WT, CCR2−/− and iCCR−/− mice. Frequencies expressed as % of CD45+, assessed via flow cytometry. ii) quantification of leukocyte numbers in the fat pad of 20 week old iCCR^−/−^ males compared to WT males, assessed via flow cytometry. **C**: Quantification of macrophage subsets (total, CD11c+ and CD11c-) in the fat pad of 20 week old iCCR−/− males compared to WT males, assessed via flow cytometry. **D**: Frequencies of CD11c+ (blue) vs CD11c- (red) adipose tissue macrophages in male WT, CCR2−/− and iCCR−/− male mice (as proportion of total CD64+ F480+ macrophages) at 8-10 weeks and at 16+ weeks. For 5B and C, Counts were normalised to the average of WT counts (age and gender matched) of each leukocyte subset to obtain values expressed as fold of WT. Unpaired t test with Welch’s correction was performed in 5A-C to determine statistical significance, with a p value of 0.05 determined as significant. *p<0.05, **p<0.01, ***p<0.001, ****p<0.0001.

### The Fat Pad Phenotype can be induced in WT females in a model of skin inflammation

Given the association of the phenotype with inflammation, we next tried to recapitulate it by inducing inflammation in WT female mice using the Imiquimod (IMQ) model of skin inflammation(16). Following IMQ treatment, WT females showed a dramatic reduction in inguinal fat pad weight and an increase in lymph node weight compared to females treated with control cream (Figure 6Ai, 6Aii). This phenotype was also apparent in IMQ-treated CCR2−/− female mice. In keeping with the fact that IMQ applied to the dorsal skin enters the circulation and thus become systemic(17), we observed a similar effect in axillary fat pad and lymph nodes (Supplementary Figure 4A) associated with an increase in plasma markers of inflammation (Supplementary Figure 5). Inguinal lymph node and fat pad weight also correlated with each other, but again the function could be divided in two: vehicle treated females appeared to have no correlation between lymph node and fat pad size, whilst IMQ-treated females showed a strong negative correlation between inguinal lymph node size and fat pad weight (Figure 6B). Analysis of the cellular content of the inguinal fat pad revealed an increase in CD45+ cells in IMQ-treated WT and CCR2−/− females (Figure 6Ci) associated with a very strong (~5 fold) increase in macrophage numbers (Figure 6Cii). Unlike fighting CCR2−/− males, we observed no increase in neutrophils, eosinophils, or lymphocyte numbers in the fat pad (Supplementary Figure 4B). Similar to CCR2−/− fighting males, fat pad macrophages in WT females treated with IMQ showed a significant increase in the levels of CD11b and CD11c (Supplementary Figure 4Ci, 4Cii).

**Figure 6.**
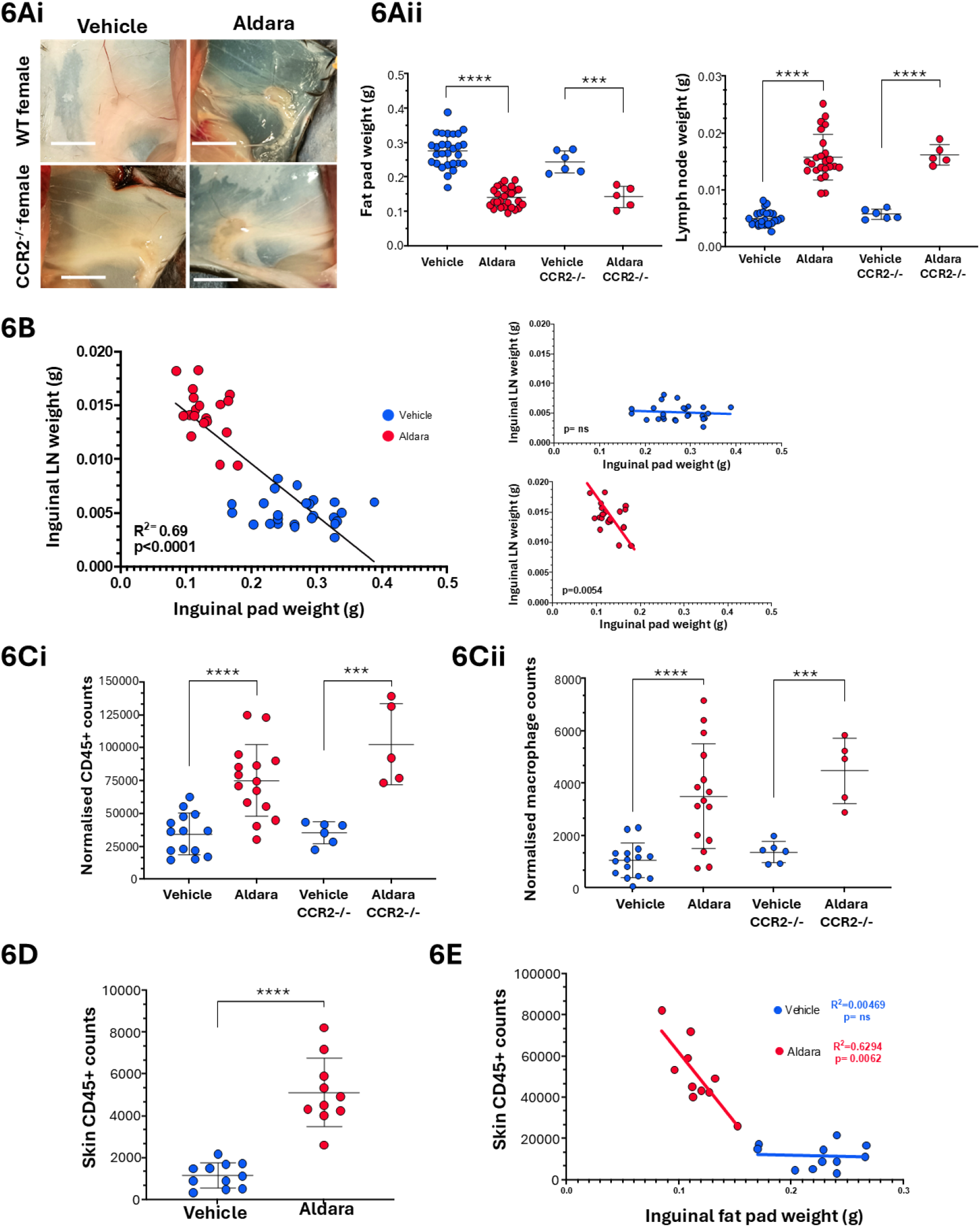
The Fat Pad Phenotype can be induced in WT females in a model of skin inflammation. **A**: i) In situ orientation of inguinal fat pad and inguinal lymph node in WT and CCR2−/− females at rest and after 3 consecutive days of Aldara application to the skin, with ii) fat pad and lymph node weight quantification. Each point represents the combined macrophage content of weight of both inguinal fat pads of a single mouse. **B**: Correlation of inguinal fat pad (x-axis) and lymph node (y-axis) weight, both expressed in grams, of WT females receiving vehicle (blue) or Aldara cream (red) to the shaved back for 3 consecutive days and culled at Day 4. Simple linear regression was significantly non-zero (p<0.0001), with an R2 values of 0.69. The vehicle only group (blue) does not show any significant correlation, while the correlation is still significant in the Aldara only group (red). **C**: i) CD45+ cell content of the inguinal fat pad after 3 consecutive days of Aldara application, assessed via flow cytometry, in WT and CCR2−/− females. Blue= vehicle control, Red= aldara cream. Leukocyte numbers were normalised to the weight of the inguinal fat sampled. Each point represents the combined inguinal fat pads of a single mouse. ii) Quantification of adipose tissue macrophages (CD64+ F480+) in the inguinal fat pad after 3 consecutive days of Aldara application, assessed via flow cytometry, in WT and CCR2−/− females. Blue= vehicle control, Red= aldara cream. Macrophage numbers were normalised to the weight of the inguinal fat sampled. Each point represents the combined inguinal fat pads of a single mouse. **D**: Quantification of live leukocytes (CD45+) in the skin of WT females after 3 consecutive days of Aldara application, assessed via flow cytometry. Blue= vehicle control, Red= aldara cream. Leukocyte numbers were normalised to the weight of the inguinal fat sampled. **E**: Correlation between inguinal fat pad weight (x-axis) and number of normalised CD45+ cells in skin (y-axis) of WT females receiving vehicle (blue) or Aldara cream (red) to the shaved back for 3 consecutive days and culled at Day 4. Simple linear regression was significantly non-zero (p<0.0062) only in females receiving Aldara, with an R^2^ values of 0.6294. The correlation was not significant in vehicle treated females (R^2^=0.00469). Unpaired t test with Welch’s correction was performed in 6A-D to determine statistical significance, with a p value of 0.05 determined as significant. ***P<0.001 ****p<0.0001

As expected, WT females treated with IMQ showed a strong increase in number of CD45+ cells in the skin (Figure 6D), and most of this increase was attributed to macrophages, however eosinophils, monocytes, DCs and lymphocytes were also significantly increased (Supplementary Figure 4D). Similar to co-housed CCR2−/− males, we observed a strong correlation between the number of CD45+ cells in the skin and the inguinal fat pad weight (Figure 6E), again suggesting that the extent of skin inflammation is correlated with local lymph node and fat pad size.

Overall, these data demonstrate that the phenotype observed in CCR2−/− male mice can be recapitulated in inflamed WT female mice and highlight macrophages as a potential driver cellular population. Our data further demonstrate, again, that the phenotype and the macrophage accumulation are independent of CCR2.

### The fat pad phenotype is at least partially dependent on macrophages

Given the common emergence of a distinct macrophage population in the CCR2−/− male inguinal fat pads, as well as in WT female mice treated with IMQ, we next assessed the transcriptome, and the potential role, of this population in inguinal fat pad, and lymph node, remodelling. To determine the transcriptional profile of the ATMs that emerge in inflamed fat pads we used ATMs from resting, or IMQ-treated, females as this inflammatory model allows for precisely timed kinetic analysis. Principal component analysis revealed that the ATM in inflamed fat pads were transcriptionally distinct from those present in resting fat pads with approximately 1100 upregulated and 700 downregulated transcripts in inflamed ATMs (Figure 7A). Bulk comparison of the control and Aldara-treated ATMs revealed that both display a core macrophage signature but that, whilst those from control female mice resembled standard inflammatory macrophages, those from Aldara-treated mice represented a distinct macrophage population which displayed upregulation of a range of genes involved in lipid metabolism, lipid homeostasis and cholesterol efflux (Figure 7B). Further analysis of the transcripts preferentially expressed in the ATM from the IMQ-treated mice revealed considerable overlap with transcripts preferentially expressed in a previously described monocyte derived population of fat pad macrophages (Figure 7C) which enter fat pads in response to either high-fat diet or calorific restriction(18). Overall, these transcriptomic data indicate that the inflammation-associated fat pad is characterised by the presence of an atypical macrophage population which displays enhanced expression of lipid-metabolising genes.

**Figure 7.**
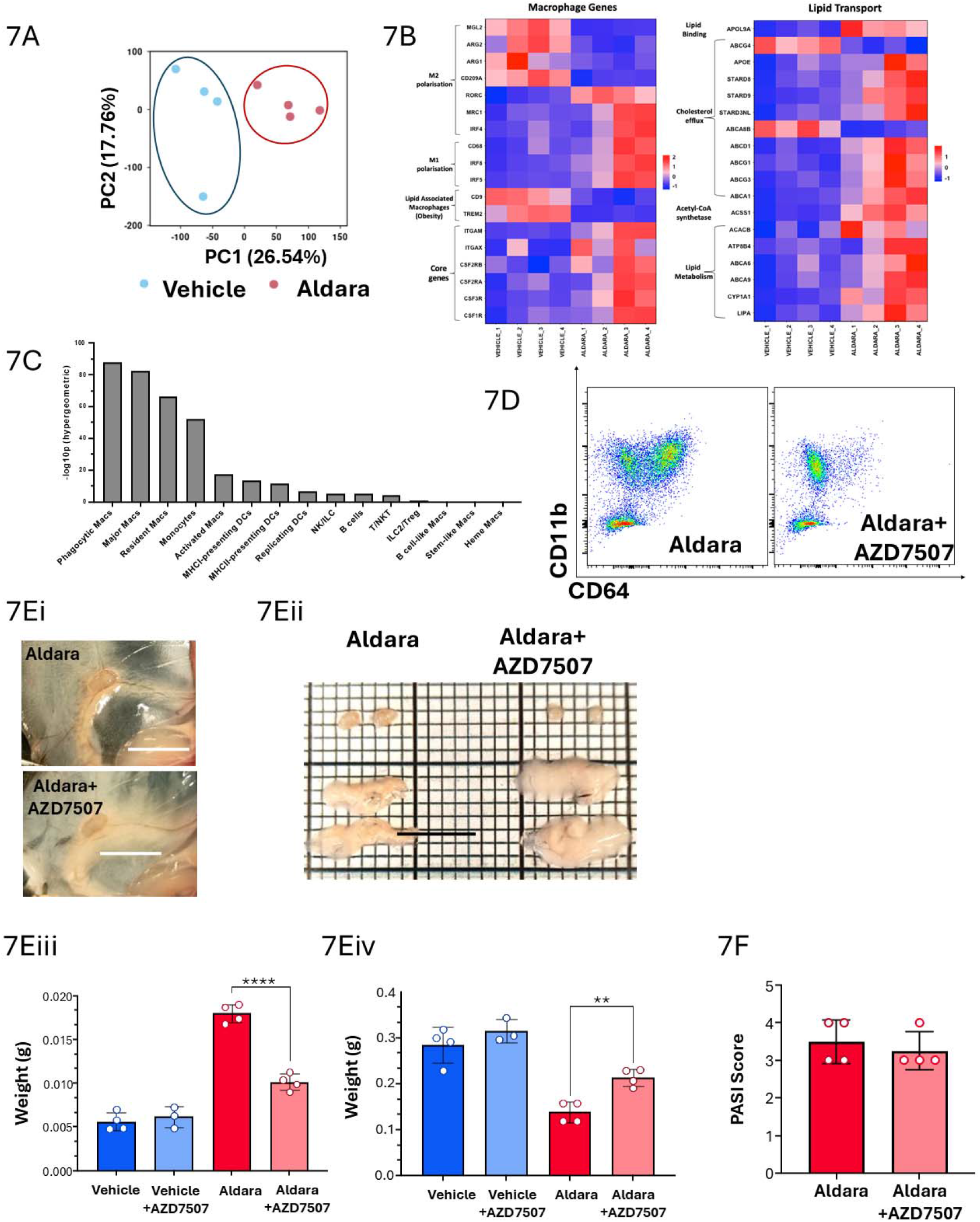
The fat pad phenotype is at least partially dependent on macrophages. **A**: Principal component analysis (PCA) plot illustrating distinct clustering of RNA samples from the inguinal fat pad of Aldara and vehicle treated female mice. Each point represents the pooled macrophage population of the inguinal fat pads of 4 mice. **B**: Heatmaps showing significantly upregulated and downregulated genes involved in macrophage function and lipid transport between inguinal fat pad macrophages from Aldara-treated or vehicle-treated mice. **C**: Overlap of the Aldara treated inguinal fat-pad macrophage RNA signature with the single cell RNA Sequencing data-set of visceral adipose tissue leukocytes from Weinstock et al, 2019. For each cell type, the markers were overlapped with the DEGs upregulated in Aldara. An overlap fold and p-value was calculated using a hypergeometric test. **D**: Flow cytometry plot showing reduction in macrophages (CD11b+CD64+) in the fat pads of AZD7507 treated mice. **E**: i) in situ fat pads and ii) excised lymph nodes from aldara, and aldara plus AZD7507, treated mice, iii) Inguinal lymph node and iv) fat pad weight of female mice before and after 3 days of Aldara induced inflammation, with and without pre-treatment with CSF1R inhibitor AZD7507. F: PASI (Psoriasis area and severity index) score of the skin of WT females and females treated with CSF1R inhibitor AZD7507, after 3 days of daily Aldara application to the skin. Unpaired t test was performed to determine statistical significance, with a p value of 0.05 determined as significant.**p<0.01, ****p<0.0001

To determine the overall contribution of macrophages to the observed phenotype, we repeated the IMQ model in female mice in the presence or absence of a pharmacological blocker of CSF1-R(19, 20). This blocker successfully reduced fat pad macrophage numbers (Figure 7D and Supplementary Figure S6) in both lymph nodes and fat pads. In addition, a small reduction in dendritic cells was seen in the fat pads. Interestingly in the lymph nodes, use of the blocker led to a comprehensive reduction in lymph node cell types. Whilst the blocker had no effect on the inguinal fat pads or lymph node in vehicle-treated animals, it ameliorated both the reduction in fat pad, and increase in lymph node, weight in the IMQ treated mice (Figure 7Ei, ii, iii and iv)). Assessment of the extent of IMQ-induced cutaneous inflammation using a modified PASI score(16) indicated, surprisingly, that CSF1-R blockade had no effect on the overall severity of the skin inflammation following IMQ treatment (Figure 7F).

Overall, these data reveal the presence of an atypical macrophage population in the remodelled fat pad and demonstrate a role for macrophages in the inguinal fat pad and lymph node remodelling downstream of inflamed skin.

### The fat pad phenotype is associated with lipolysis and free fatty acid availability

A number of genes with key roles in fatty acid metabolism were up regulated in the inflamed fat pad ATMs (Figure 8A). This is indicative of active lipolysis in the fat pads and, given the importance of fatty acids as energy sources for the immune system(8, 10, 21, 22), suggested that the fat pad may act as a source of energetics for the embedded lymph node. This suggestion is further supported by an analysis showing the close temporal association of reduction in fat pad weight and increase in lymph node weight (Figure 8B). We therefore next examined free fatty acid levels in the fat pads and the associated lymph nodes. As shown in the exemplary data in Figure 8C, the availability of a number of free fatty acid species demonstrated significant increases in both the fat pad, and lymph nodes, over a time-course covering days 1, 3 and 5 post-IMQ treatment. Notably the levels of the individual fatty acids generally increased in the fat pads prior to the increase in the lymph nodes. In addition, at least for the C20 fatty acids, levels in the fat pads far exceed those in the lymph nodes. Together these observations suggest that the fatty acids have originated from the fat pads and not the lymph nodes and raise the possibility of fatty acid exchange from the lipolysis-active fat pads to the lymph nodes.

**Figure 8.**
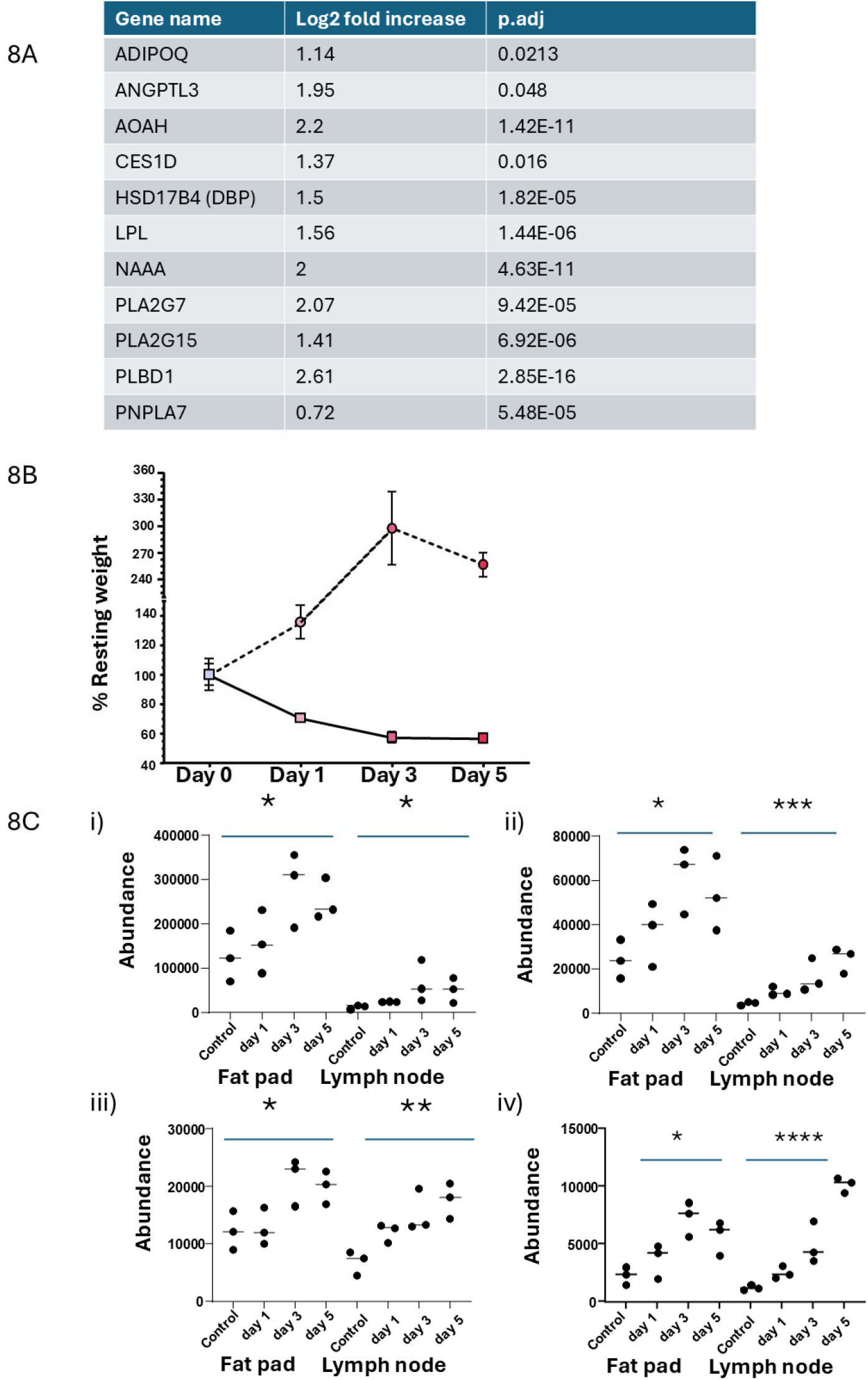
The fat pad phenotype is associated with lipolysis and free fatty acid availability. **A**: List of transcripts encoding proteins involved in lipolysis which are upregulated in the adipose tissue macrophages in Aldara treated mice. **B**: Plot showing the reciprocal relationship between reduction in fat pad and lymph node weight over the time course of Aldara treatment. **C**: Exemplar free fatty acid analysis showing levels of i) FA 20:1; ii) FA 20:2; iii) FA 22:1; iv) FA 22:2. Statistics were analysed using one-way ANOVA with a p-value of 0.05 determined as significant. *p<0.05, **p<0.01, ***p<0.001, ****p<0.0001.

## DISCUSSION

It has been realised for some time that the high energy requirements of an immune response are met, in significant part, by lipolysis of perinodal fat deposits(21, 22). However, the trigger for this has remained elusive. Here we demonstrate a clear link between the extent of peripheral cutaneous inflammation and remodelling of draining lymph nodes and perinodal fat pads. Specifically, we see a rapid reduction in the size of the fat pads with a reciprocal increase in the size of the embedded lymph node. This phenotype was seen downstream of fighting wounds, in CCR2−/− male mice, and in WT female mice treated with the inflammatory TLR7 agonist IMQ and was associated with the accumulation of a specific macrophage population in the atrophied fat pads. Notably, this phenotype was observed in both the fighting CCR2−/− males, as well as IMQ treated CCR2−/− females, demonstrating that the phenotype is independent of CCR2 at the peripheral inflamed site and the draining fat pad and lymph node.

Interestingly, we observed a more robust inflammatory response to fighting-associated wounding in co-housed CCR2−/− male mice compared to isolated CCR2−/− male mice or WT male mice. This is a counterintuitive observation given the non-redundant role for CCR2, and its ligands, in monocyte egress from the bone marrow and recruitment to inflamed sites(11, 12, 23). Notably, similar results have been reported from other studies on inflammation in CCR2−/− mice(24-27) and potentially reflect the inability to recruit a specific CCR2-dependent anti-inflammatory macrophage population which would otherwise restrict the extent of the inflammatory response(27). It is also important to note that, despite the anticipated monocytopenia that is seen in CCR2−/− mice, there is no difference in the total number of macrophages in the inflamed skin of co-housed and WT male mice suggesting that in the CCR2−/− mice the macrophages are derived from CCR2-independent monocyte recruitment or in situ macrophage proliferation.

In terms of the altered macrophage population in the atrophied perinodal fat pad, transcript omics analysis reveals these to be similar to a population of ATM described by Weinstock and colleagues as being markedly increased upon calorific restriction(18). There are many other reports of macrophage recruitment to perinodal fat pads following inflammatory stress or calorific restriction(28, 29). The ATMs identified in the current study are characterised by relatively increased expression of genes involved in lipolysis and analysis of free fatty acids indicate active lipolysis in the fat pad in response to inflammation. Furthermore, the use of a well-characterised pharmacological blocker of CSF1R(20) clearly indicates a requirement for macrophages in the remodelling of the perinodal fat pad and lymph nodes. Overall, this points to the recruitment (or in situ proliferation) of a CCR2- independent macrophage population with enhanced lipolytic activity. Interestingly, there have been reports of a direct conduit system between perinodal fat deposits and the embedded lymph nodes(30) suggesting a clear route for transfer of fatty acids from the fat pad to the lymph node. It is important to note that, whilst our data is compatible with a model of fatty acid movement from fat pad to lymph node, it remains possible that the linked alteration in fat pads and lymph node could be the result of other phenomena such as lymph node intrinsic metabolism or plasma spillover of fatty acids.

In summary, our study demonstrates a striking link between peripheral inflammation and perinodal fat pad atrophy and lymph node expansion. This suggests that inflammation is a key determinant of local energy provision from perinodal fat pads to lymph nodes and that this can be seen independently of antigenic challenge. Our data are also important in that they suggest that pro inflammatory vaccine adjuvants may mediate aspects of their function(31, 32), remote from the injection site, by priming downstream fat pads and embedded lymph nodes for the development of subsequent immune responses.

## Acknowledgements

The authors acknowledge the help of the MVLS Core Cellular Analysis Facility. This work was funded by the Wellcome Trust (217093/Z/19/Z) and the Medical Research Council (MR/V010972/1).

## Materials and Methods

### Mice and co-housing

Wild-type (WT), CCR2−/− and iCCR−/− mice are on a C57BL/6 background, and were maintained as previously described(11, 23), in a specific-pathogen free facility at the University of Glasgow under the auspices of a U.K Home office Licence. All animal procedures were approved by the local ethical review committee. Animals used were both male and female, ‘young’ mice were classified as 8-10 weeks of age, adult mice were 14-16 weeks of age, while ‘old’ mice were classified as 20+ weeks of age. To assess the impact of fighting and the contribution of skin injuries to the inflammatory phenotype observed, 8-10 week old male litters were separated and co-housed with individuals from other litters of the same age in groups of 4-5 for 6 weeks.

### Adipose Tissue, Skin and Lymph Node digestion

Mice were sacrificed and tissues extracted for flow cytometry and RNA sequencing. Excised adipose tissue (inguinal fat pads, infrascapular brown fat, visceral white fat and axillary fat pads) were coarsely cut and digested for 1 hour on a Thermoshaker at 37°C at 850rpm in 1ml of Liberase in Hanks buffered saline (with Ca and Mg, ThermoFisher Scientific), at a final concentration of 0.44 Wunch units/sample. Samples were vortexed at the 30-minute mark, placed on ice after 1 hour, and filtered through a 100µm mesh strainer in a 50ml falcon, and the filter was washed with 10ml of PBS. Samples were spun down at 300g for 5 minutes, and the pellet was resuspended and passed through a 70µm mesh strainer. Skin samples were processed in a similar way however digestion was performed at 1000rpm, with vortexing every 15 minutes to encourage dissociation. Lymph nodes were collected and mechanically digested through a 70µm mesh strainer with the aid of a 2ml syringe plunger. The mesh was washed with 10ml PBS, and the sample was spun down at 300g for 5 minutes. The resulting pellet was resuspended in 10ml of PBS and re-filtered through a new 70um strainer to remove remaining fat and connective tissue.

### Isolation of Adipose tissue macrophages

Cell counts from digested and filtered adipose tissue samples were obtained via a Luna Cell Count Reader, and the appropriate amount of Anti-F480 MicroBeads Ultrapure (Miltenyi Biotech) were added to each sample to magnetically isolated adipose tissue macrophages according to the manufacturer’s protocol. Briefly, 10 µl of Anti-F480 MicroBeads Ultrapure were added for every 10^7^ cells, and incubation was performed for 15 minutes on ice in the dark. Cells were washed with 10ml of FACS Buffer and centrifuged at 300g for 10 minutes. The pellet was resuspended in 500µl, and the suspension was placed on an MS MACS column on an OctoMacs separator (Miltenyi Biotech). The flow-through containing unlabelled cells was discarded, while the eluted fraction containing magnetically labelled cells was collected and enriched over a second MS columns to achieve higher purities. At the end of the second magnetic isolation, a small portion of each sample was collected and stained with CD64, Ly6G and SiglecF fluorescent antibodies to test macrophage content and amounts contaminating cells via flow cytometry. RNA extraction was performed on samples with >90% CD64+ cells.

### Flow cytometry

Starting from single-cell suspensions, cells were first stained with a fixable viability dye (Zombie Aqua; BioLegend), and Fc receptors were blocked using FcR Blocking-Reagent (Miltenyi Biotec). Surface marker staining was performed with the appropriate Ab mix diluted in FACS Buffer (PBS/2 mM/EDTA/0.5% FBS) on ice for 20 min. Cells were washed, fixed with Fixation Buffer (420801; BioLegend), and resuspended in FACS buffer before analysis. Data were acquired using either the Fortessa (BD Biosciences) flow cytometer and analyzed using the FlowJo 10 software. Voltages and compensation were determined using UltraComp eBeads (01-2222-42; Bioscience) as single controls. Positive staining was determined using Fluorescence-Minus-One controls. The gating strategy and antibody list for inguinal fat pad leukocytes are outlined in Supplementary figure 7.

### AZD7507 Treatment

To achieve macrophage depletion, mice received 100μl of AZD7507(20) (MedChemExpress, HY-117244), an orally active CSF-1R inhibitor, at a concentration of 100mg/kg (25mg/mL). 250mg of AZD7507 were resuspended in 10ml of MilliQ water with 0.5% HPMC and 0.1% tween 80. 100µl of AZD7507 suspension was added to 1ml syringes, and administered to mice via oral gavage, twice a day, for 8 days. Mouse weights were monitored daily, and after 8 days, the back of the mice was shaved to allow for Aldara application.

### Aldara/Imiquimod Model

WT and CCR2−/− female mice were each treated with a ~62.5 mg dose of Aldara™ (Meda AB) cream containing 5% Imiquimod or an equivalent volume of control cream (Boots Aqueous Cream/10% Vaseline Lanette Cream) by application to shaved dorsal skin near the base of the tail (~3mm^2^ area)(16). Cream was applied for 3 consecutive days (day 1, day 2 and day 3) and the mice culled 24h after the final application. Mice were monitored daily for weight loss. Following euthanasia blood was collected immediately using a syringe flushed with 0.5 M EDTA. Following blood collection, mice were transcardially perfused with 20ml of ice-cold PBS. When tissue was required for immunostaining, mice were subsequently perfused with 4% PFA. Development of a psoriasiform pathology was carried out as described(16). Inflammatory scoring was performed by an independent investigator blinded to treatment allocation. Formal randomization was not implemented because AZD-treated animals required additional gavage treatment and associated experimental handling.

### RNA Extraction and processing

After F480-magnetic positive selection, the eluted cell suspensions from 6 inguinal fat pads were pooled together to obtain enough ATMs (around 30,000 cells) for low-input RNA sequencing. Cells were spun down at 1000g for 10 minutes, lysed, and spun again to remove bound magnetic beads before proceeding with RNA extraction. RNA was extracted using the RNeasy Microkit with on-column DNAse I digestion (Qiagen), following manufacturer’s instructions, obtaining samples of around 100-150ng of total RNA in water.

RNA samples were then sent to a commercial sequencing provider (Novogene), for RNA QC, mRNA library preparation (poly-A enrichment) and Illumina Sequencing PE150.and sequencing. Briefly, raw reads of fastq format were processed through in-house scripts, removing adapter, ploy-N and low-quality reads from raw data. Index of the reference genome was built and paired-end clean 1 reads were aligned using Hisat2 v2.0.5. FeatureCounts v1.5.0-p3 was used to count the reads for each gene. Differential expression analysis of two conditions/groups was performed using the DESeq2Rpackage (1.20.0). The resulting p-values were adjusted suing the Benjamini and Hochberg’s approach for controlling the false discovery rate. Genes with an adjusted P-value <= 0.05 found by DESeq2 were assigned as differentially expressed

#### Comparison of scRNA cell type expression with Aldara

The Weinstock et al dataset(18) was downloaded from GEO (GSE141036) as a Seurat object. The data was analysed and converted into pseudo bulk. In detail, Seurat (v5.0.3)(ref - 2) was used to load the dataset. Cells were filtered for nFeature_RNA > 200 & percent_mito < 5. The data was SCT transformed, and PCA, find clusters, and generate UMAP performed using PC 1-30. Cluster markers were identified using FindAllMarkers with the settings only.pos = TRUE, min.pct = 0.25, logfc.threshold = 0.25. All other settings were left to default. As expected the resulting UMAP and clusters were highly similar to the manuscript, and so cell types were matched. Next a per cell type pseudo-bulk count table was generated from the raw counts, using a loop in R. This was normalised into an expression table using DESeq2 (v3.18)(ref - 3), under default settings. To compare to the Aldara dataset the mean value for each gene in control and Aldara groups was calculated. Both the Aldara and pseudo-bulk data were filtered to include only the marker genes from the overlap analysis (see above). Finally, the pseudo- bulk expression values for each cell type were compared to the equivalent values in the control or Aldara group using a Spearman correlation.

### Free fatty acid analysis

A methanol extraction of foetal bovine serum was used as a quality control sample and used for instrument conditioning prior to sample injection. Methanol extracts from samples were centrifuged at 13000 rpm for 5 min at 4°C, to separate protein precipitate from the methanol supernatant. For each sample, an aliquot of supernatant was diluted 1:1 (v/v) with MilliQ water (Merck Millipore) spiked with C17:0 fatty acid at 0.8 µg/ml and transferred into an autosampler vial. Sample vials were kept at 8°C in an Acquity Premier FTN autosampler (Waters corporation). An aliquot of 10µL of each sample was injected onto CORTECS UPLC T3 1.6 µm, 3 x 100mm column (Waters corporation) held at 50°C within an Acquity Premier column manager (Waters corporation). Free fatty acids were separated using Acquity Premier BSM UPLC pump (Waters Corporation), using 0.2 mM Ammonium formate with 0.01% formic acid for solvent A, and Isopropanol:Acetonitrile (1:1 v/v) 0.2 mM ammonium formate with 0.01% formic acid for solvent B. Elution gradient was as follows: 30% B for 0.1 min, 30-65% in 0.9 min, 65-75% B in 2.5 min, 75-98% B for 1.5 min, 98% held for 1min before re-equilibration at 30% for 2 minutes. Fatty acids were detected in negative ion mode using a Xevo TQ Absolute (Waters Corporation) triple quadrupole mass spectrometer. Ionisation conditions were heated electrospray source with capillary voltage 1.9kV, desolvation temperature 500°C, desolvation gas 1000L/Hr, cone gas 150L/Hr and source temperature of 120°C. Fatty acid ions were detected using MRM mode with parent to parent transitions (see attached MRM table), with a +/− 0.5 min window around the expected elution time, with a cone voltage of 30V and collision energy of 15eV. 41 transitions were acquired from C6:0 to C24:0, including common polyunsaturated fatty acids. The following fatty acids were matched by mass and retention time to authentic standards, C6:0, C8:0, C10:0, C12:0, C14:0, C16:0, C18:0, C20:0, C22:0, C24:0, alpha-linolenic acid (18:3n3), gamma-Linolenic acid (18:3n6) and Arachidonic acid (C20:4n6). All other reported fatty acids were determined based on formula mass and estimated retention time. Peak picking of the acquired data was performed using TargetLynx (Waters Corporation), before exporting to a text file. The results text file was then read using custom scripts in R (version 4.41). Signals for background ions were averaged and then subtracted from analyte signals. Any negative values were then removed from the analysis. Chains shorter than C12:0 were determined to be too variable, along with the internal standard C17, which was spiked into the sample, and therefore not reported. The resulting data table was then saved as an Excel spreadsheet.

### Statistical analysis

All statistical analyses were performed using Graph Pad Prism

