## Supplemental figures for "Macrophages drive inguinal fat pad and lymph node remodelling in response to peripheral inflammation"

**Supplementary data. Bartolini et al.**

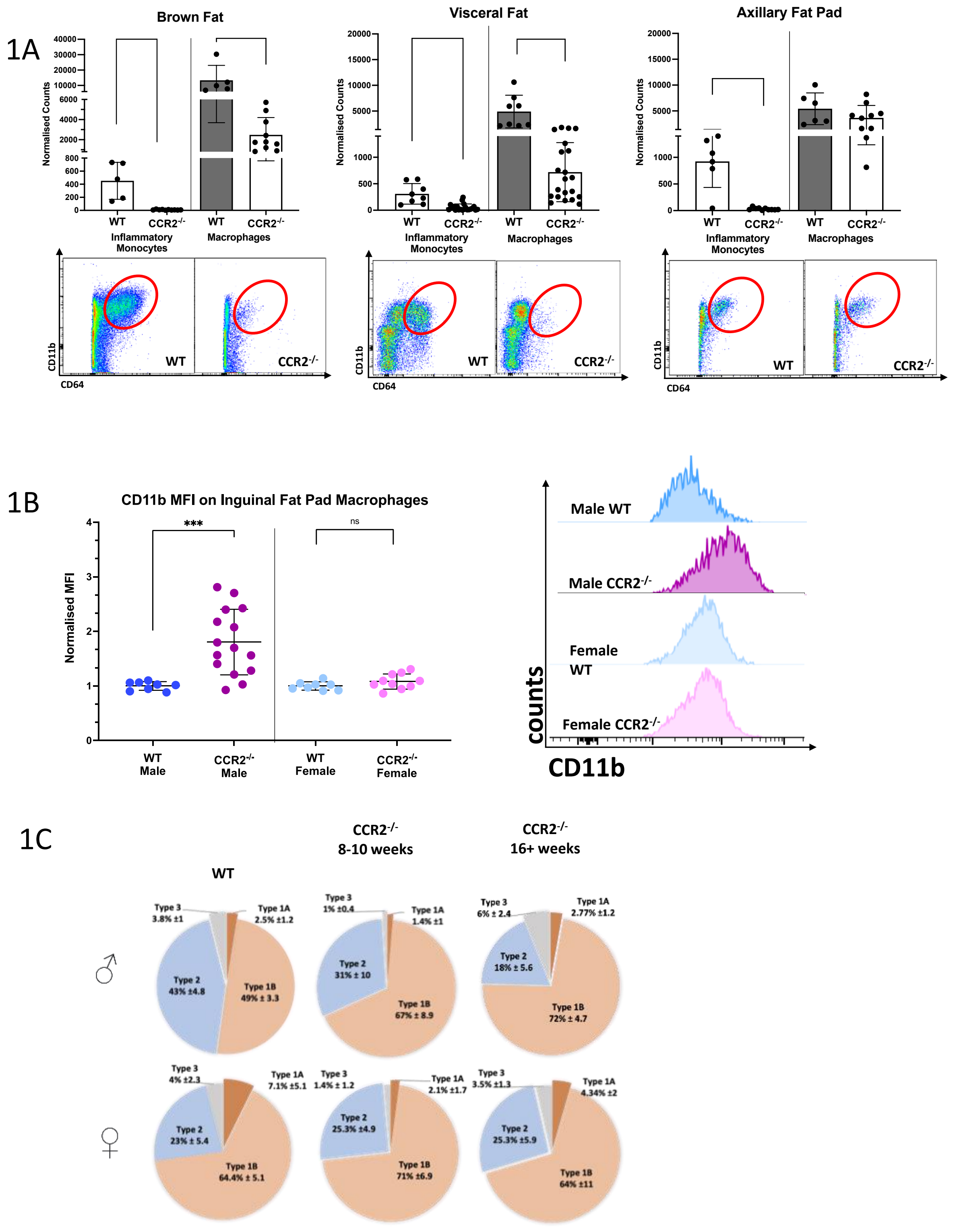


**Supplementary 1**

**1A**- Counts of inflammatory monocytes (CD11b+Ly6C++) and macrophages (CD11b+CD64+) in brown fat, visceral fat, and axillary fat of WT and CCR2-/- male mice, with representative FACS plots for each showing CD11b+CD64+ cells (red oval). Counts were normalised to the weight of the fat sample.

**1B**- CD11b expression measured as mean fluorescent intensity (MFI) of inguinal fat pad macrophages from WT (blue) and CCR2-/- males (purple), and WT (light blue) and CCR2-/- females (pink), with associated representative histograms.

**1C-** Frequencies of Type 1a (CD11c+ CD206-, brown), Type 1(CD11c+ CD206+, orange), Type 2 (CD11c- CD206+, blue) and Type 3 (CD11c-CD206-, grey) inguinal adipose tissue macrophages in male and female WT and CCR2-/- (as proportion of total CD64+ F480+ macrophages) at 8-10 weeks and at 16+ weeks.

Unpaired t test with Welch's correction was performed in S1A,E to determine statistical significance, with a p value of 0.05 determined as significant. ***p<0.001

2A

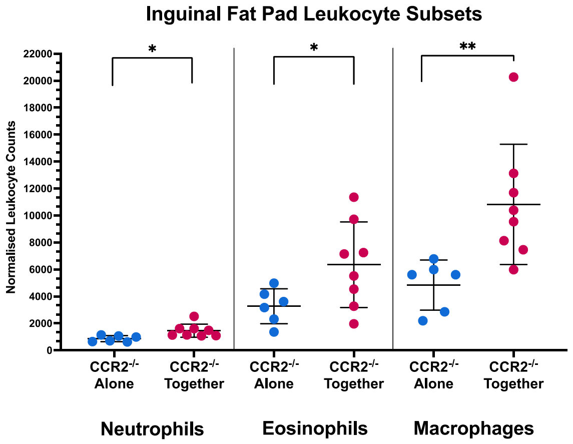


2B

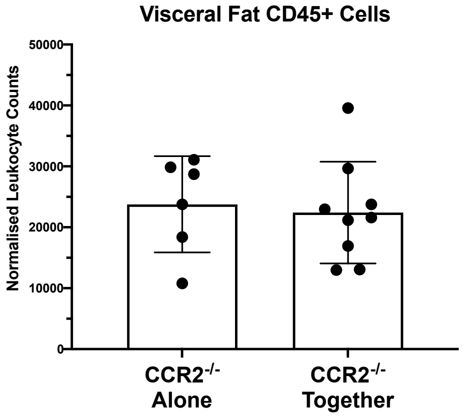


2C

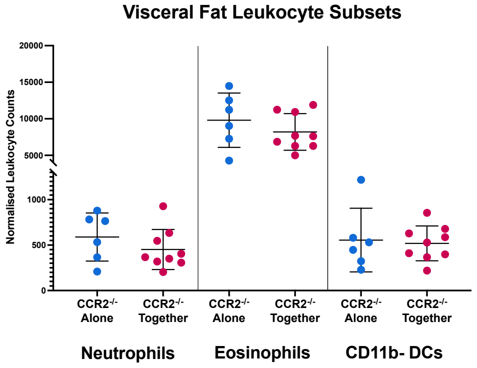


2D

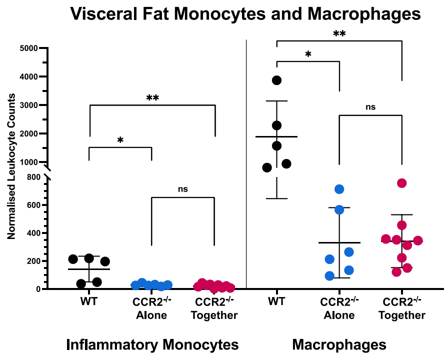


**Supplementary 2**

**2A**- Neutrophil, Eosinophil and macrophage numbers in the inguinal fat pad of CCR2-/- male mice, housed either alone or in groups of 5 for 8 weeks.
**2B**- CD45+ cell counts in visceral fat of CCR2-/- male mice, housed either alone or in groups of 5 for 8 weeks, assessed via flow cytometry and normalised to weight of visceral fat samples.
**2C**- Neutrophil, eosinophil, DC, **2D-** monocyte and macrophage numbers in the visceral fat of $CCR2^{-/-}$ male mice, housed either alone or in groups of 5 for 8 weeks.
Unpaired t test was performed to determine statistical significance. *p<0.05, **p<0.01.

3A

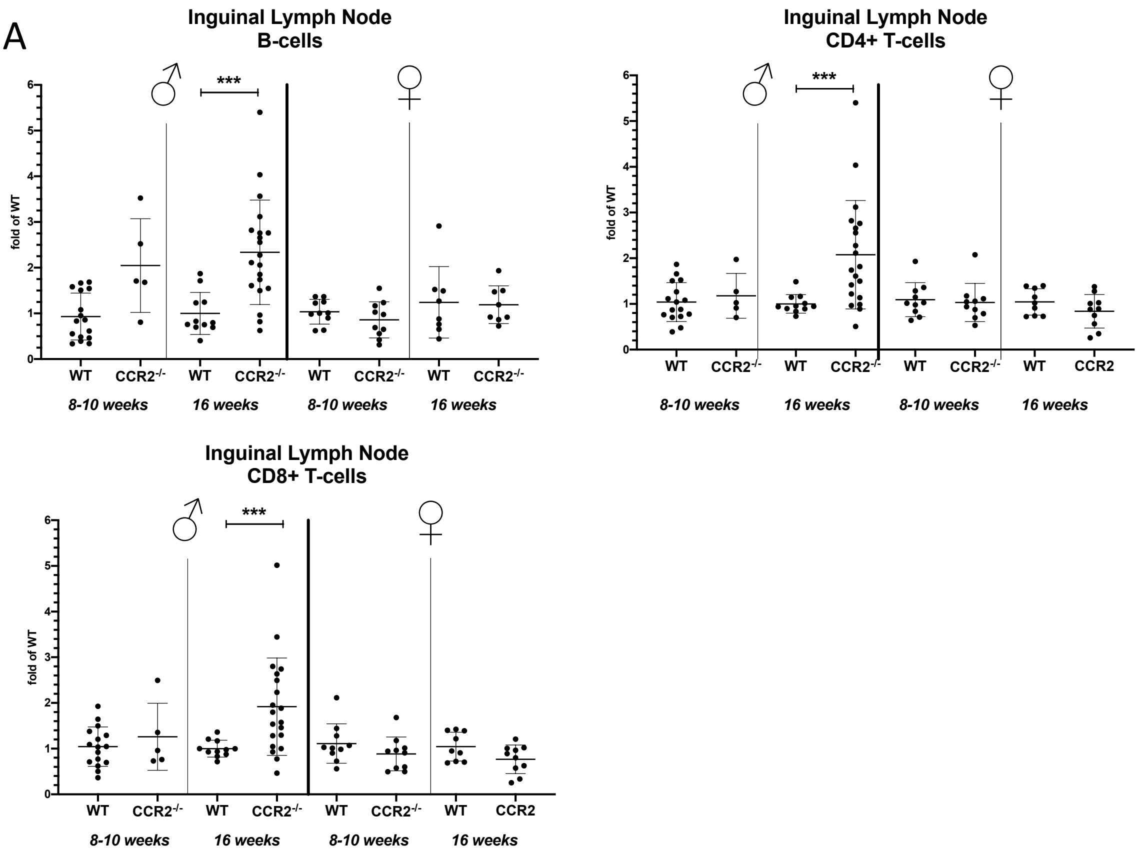


3B

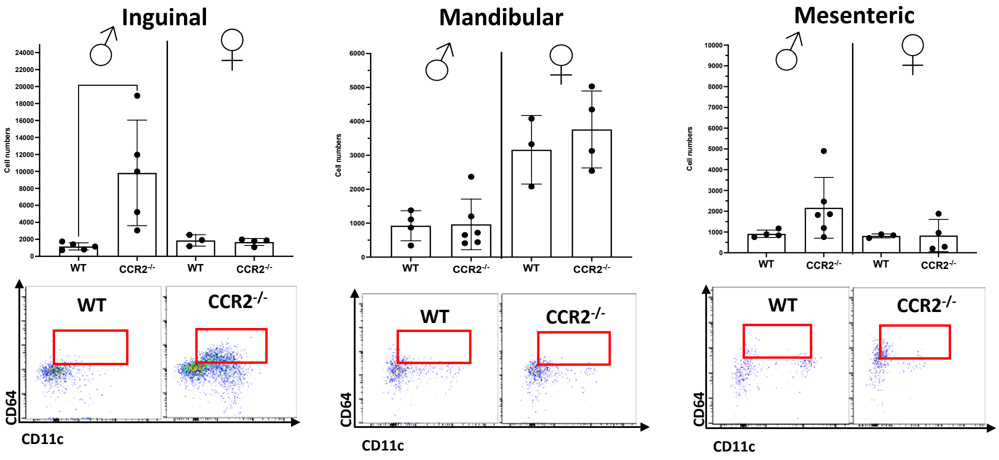


### Supplementary 3

**3A**- B-cells, CD4+ and CD8+ T-cell numbers, in WT and CCR2-/- inguinal lymph nodes, (male and female, 8-10 and 16+ weeks old) expressed as fold of 8-10 week old gender matched WT counts.

**3B**- CD11c+CD64+ Macrophage counts, assessed via flow cytometry, in inguinal, mandibular and mesenteric lymph nodes of WT and CCR2-/- mice, both male and female, with associated representative FACS plots for each, highlighting CD64+CD11c+ cells (red box).

Unpaired t test with Welch's correction was performed in S1A,E to determine statistical significance, with a p value of 0.05 determined as significant. ***p<0.001

4A

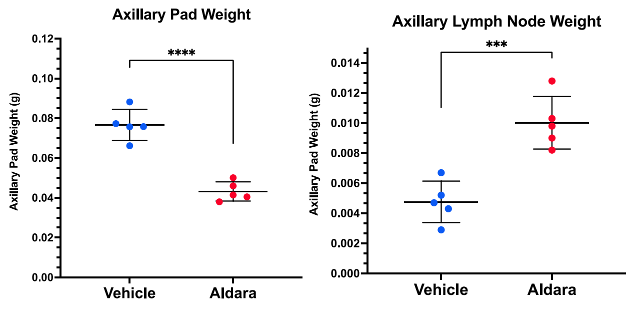


4B

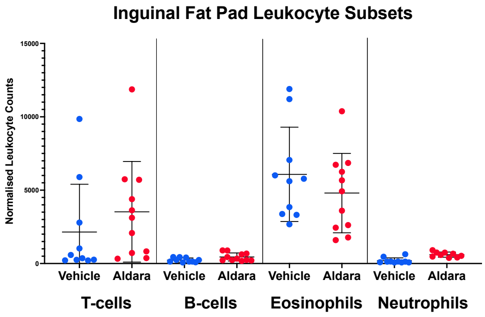


4Ci

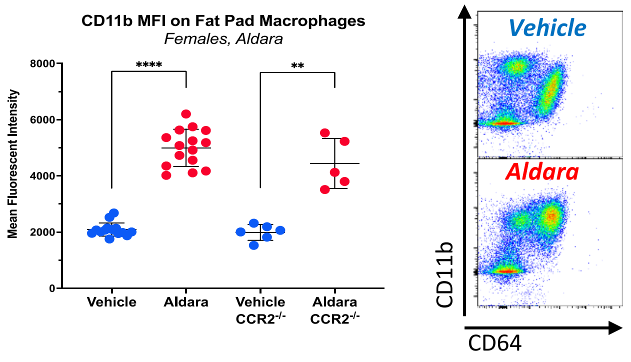


4Cii

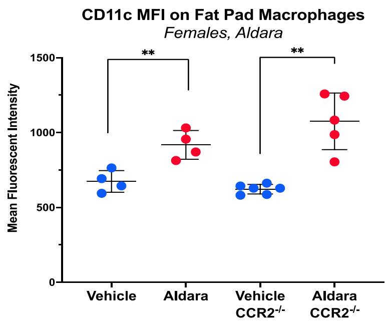


4D

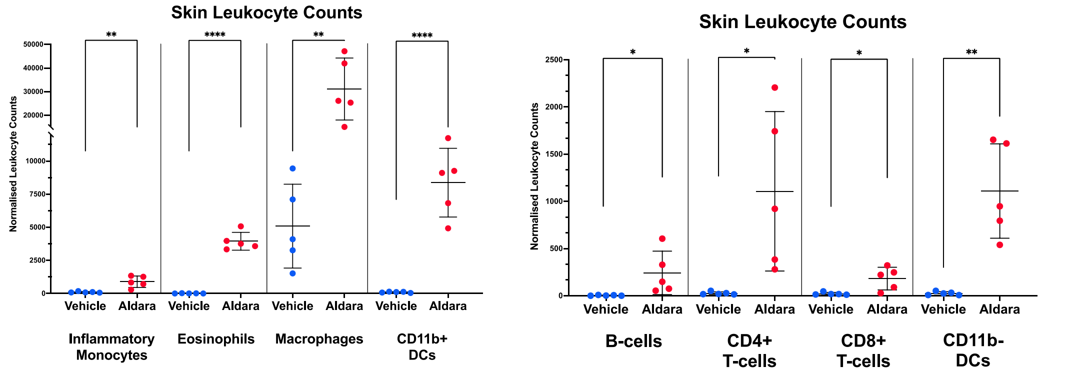


**Supplementary 4**

**4A**- Axillary fat pad and lymph node weight, in grams, of WT females treated for 3 consecutive days with Aldara (red) or vehicle (blue) cream.
**4B**- Inguinal fat pad leukocyte (T-cell, B-cell, eosinophil and neutrophil) counts in WT females treated for 3 consecutive days with Aldara (red) or vehicle (blue) cream.
**4C**- i) CD11b and ii) CD11c expression, measured as mean fluorescent intensity (MFI), of in inguinal fat pad macrophages from vehicle treated (blue) and Aldara treated (red) females, with associated representative FACS plots show the shift in CD11b expression of the CD11b+CD64+ macrophages (y-axis).

**4D**- Leukocyte (myeloid left, lymphoid right) numbers in the skin of vehicle treated (blue) and Aldara treated (red) females. Counts were normalised to the weight of the skin sampled. Unpaired t test with Welch's correction was performed to determine statistical significance. *p<0.05, **p<0.01, ***p<0.001, ****p<0.0001

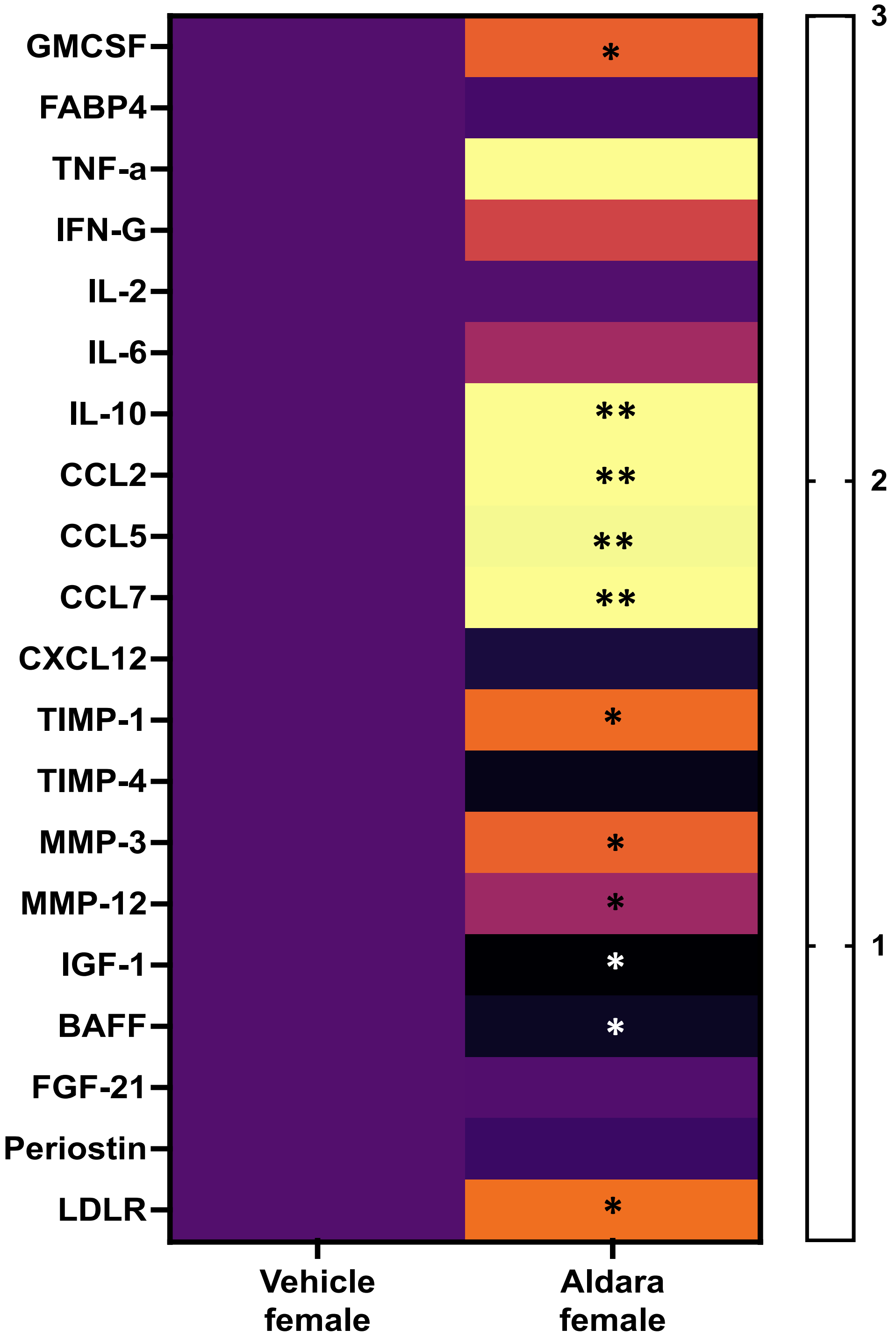


**Supplementary 5**

Plasma cytokine profiling of Aldara (n=13) and vehicle treated females (n=9) at Day 4. Data for all cytokines in heatmap expressed as 'fold of Vehicle female' concentration. Statistical comparisons between groups were performed using a two-tailed Mann–Whitney U test, significant cytokines are highlighted with an '*'.

6A

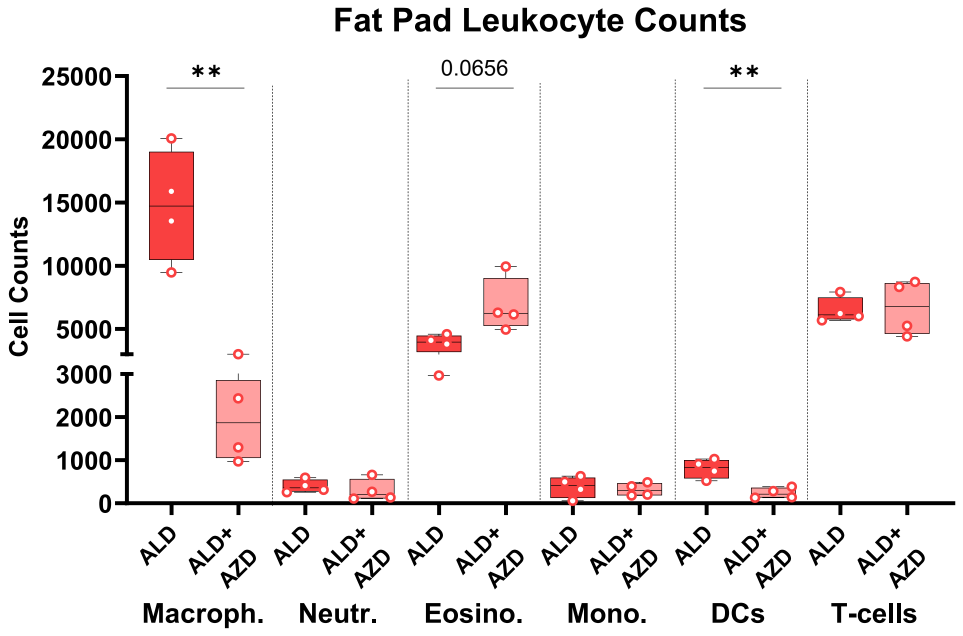


6B

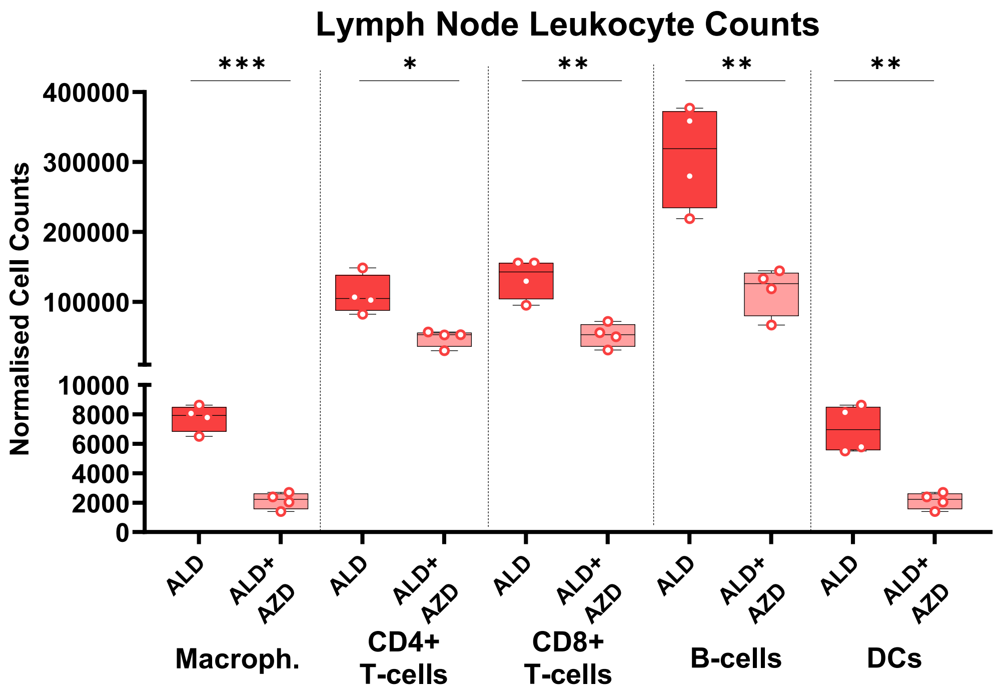


**Supplementary 6**

**6A-** normalised fat pad leukocyte counts from Aldara and Aldara plus CSF1R blocker treated mice (3 days).

**6B-** normalised lymph node leukocyte counts from Aldara and Aldara plus CSF1R blocker treated mice (3 days).

Unpaired t test with Welch's correction was performed to determine statistical significance. *$p<0.05$, **$p<0.01$, ***$p<0.001$

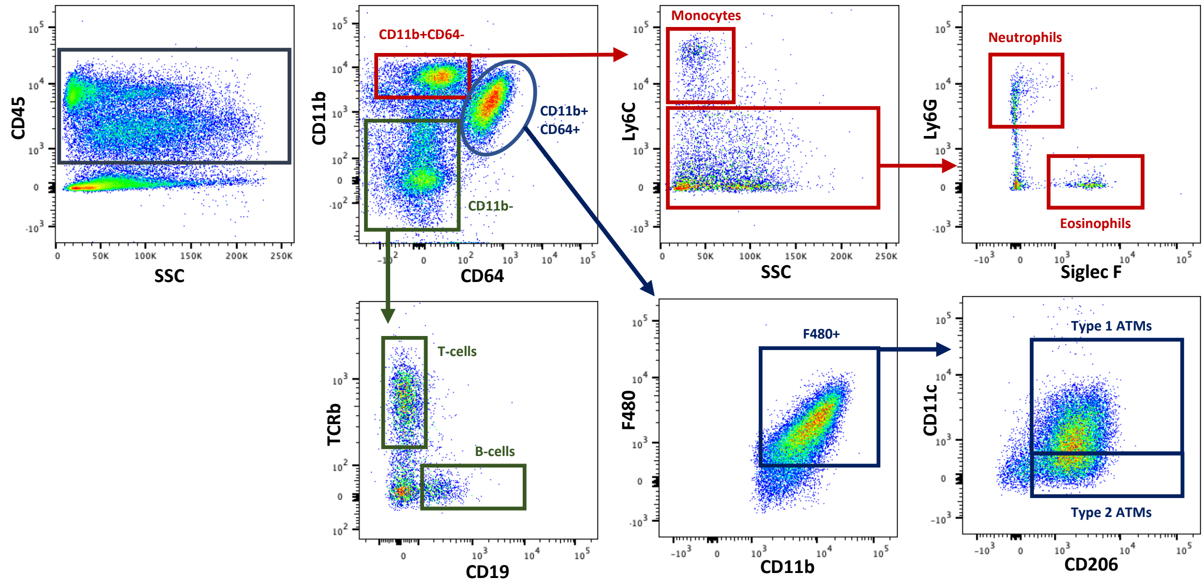


| Marker | Fluorophore | Supplier | Clone |
|---|---|---|---|
| F480 | BUV805 | BD Biosciences | T45-2342 |
| CD45 | BUV395 | BD Biosciences | HI30 |
| MHCII | BV786 | BD Biosciences | M5/114 |
| TCRβ | BV605 | Biolegend | H57-597 |
| live | BV500 | Biolegend | Zombie Aqua |
| CD64 | BV421 | Biolegend | X54-5/7.1 |
| Ly6G | FITC | Biolegend | RB6-8CC5 |
| SiglecF | PE | Biolegend | S17007L |
| CD11c | PerCP | Biolegend | N418 |
| Ly6C | Pe-Cy7 | Biolegend | HK1.4 |
| CD19 | APC | Biolegend | SJ25C1 |
| CD206 | AF700 | Biolegend | C068C2 |
| CD11b | APC-Cy7 | Biolegend | ICRF44 |

**Supplementary 7**

Gating strategy for identification of inguinal fat pad macrophages and other main leukocyte subsets. Myeloid cells including neutrophils, eosinophils and monocyte in red, lymphoid cells (B-cells and T-cells) in green, and ATMs in blue. ATMs are divided according to their expression of CD11c and CD206.
